# Transcranial Random Noise Stimulation acutely lowers the response threshold of human motor circuits

**DOI:** 10.1101/2020.10.07.329813

**Authors:** Weronika Potok, Marc Bächinger, Onno van der Groen, Andreea Loredana Cretu, Nicole Wenderoth

## Abstract

Transcranial random noise stimulation (tRNS) over cortical areas has been shown to acutely improve performance in sensory detection tasks. One explanation for this behavioural effect is stochastic resonance, a mechanism that explains how signal processing in non-linear systems can benefit from added noise. While acute noise benefits of electrical random noise stimulation have been demonstrated at the behavioural level as well as in *in vitro* preparations of neural tissue, it is currently largely unknown whether similar effects can be shown at the neural population level using neurophysiological readouts of human cortex. Here we hypothesized that acute tRNS will increase the responsiveness of primary motor cortex (M1) when probed with transcranial magnetic stimulation. Neural responsiveness was operationalized via the well-known concept of the resting motor threshold (RMT). We showed that tRNS acutely decreases RMT. This effect was small, but it was consistently replicated across four experiments including different cohorts (total N=81, 46 females, 35 males), two tRNS electrode montages, and different control conditions. Our experiments provide critical neurophysiological evidence that tRNS can acutely generate noise benefits by enhancing the neural population response of human M1.

**Significance statement:** A hallmark feature of stochastic resonance is that signal processing can benefit from added noise. This has mainly been demonstrated at the single-cell level *in vitro* where the neural response to weak input signals can be enhanced by simultaneously applying random noise. Our finding that tRNS acutely increases the excitability of corticomotor circuits extends the principle of noise benefits to the neural population level in human cortex. Our finding is in line with the notion that tRNS might affect cortical processing via the stochastic resonance phenomenon. It suggests that enhancing the response of cortical populations to an external stimulus might be one neurophysiological mechanism mediating performance improvements when tRNS is applied to sensory cortex during perception tasks.

## Introduction

Transcranial random noise stimulation (tRNS) is a non-invasive electrical brain stimulation technique whereby currents are randomly drawn from a predefined range of intensities and frequencies (Antal and Herrmann 2016). Until now most studies applied tRNS for several minutes over primary motor cortex (M1), which typically leads to an increase in corticomotor excitability relative to baseline for up to 60 min after stimulation (Terney et al. 2008; Chaieb et al. 2011, 2015; Abe et al. 2019; Moret et al. 2019) with occasional reports suggesting inhibitory effects for low intensities (Moliadze et al. 2012). The exact mechanism causing this temporary facilitation of cortical activity is unknown, however, it has been hypothesized to reflect neuroplastic changes (Terney et al. 2008).

By contrast, acute (i.e., online) effects of tRNS have been studied much less. One general hypothesis is that the brain responds to electrical noise according to a stochastic resonance (SR) phenomenon (Terney et al. 2008; Miniussi et al. 2013; van der Groen and Wenderoth 2016; van der Groen et al. 2018, 2019; Pavan et al. 2019). SR is a general mechanism that enhances the response of nonlinear systems to weak subthreshold signals by adding an optimal level of random noise (Gingl et al. 1995; McDonnell and Abbott 2009). One key indicator of the SR phenomenon in a broad sense (McDonnell and Abbott 2009) is that the investigated system “benefits” from noise, which usually refers to better detection, transmission or processing of the input signal than when no noise is present. In humans, SR effects have been mainly demonstrated via behavioural signal detection tasks whereby noise was added to the periphery. For example, the detection of low-contrast visual stimuli was significantly enhanced when the stimuli were superimposed with visual noise (Simonotto et al. 1997). Recently, a similar enhancement of visual perception has been reported when noise was directly added to visual cortex via tRNS, which improved the detection of low contrast visual stimuli (van der Groen and Wenderoth 2016), visual decision making (Ghin et al. 2018; Pavan et al. 2019; van der Groen et al. 2019), binocular rivalry (van der Groen et al. 2018), and visual training in healthy participants (Fertonani et al. 2011; Pirulli et al. 2013) and patients (Moret et al. 2018; Herpich et al. 2019). However, until now the beneficial effect of adding external electrical random noise to neural activity has mainly been studied via behavioural outcome measures in humans or via physiological single cell studies in animals (Onorato et al. 2016; Remedios et al. 2019). By contrast, it is largely unknown whether tRNS causes acute benefits when applied in-vivo to neural populations within the cortex. Here we seek to answer this question by delivering electrical noise transcranially to human primary motor cortex (M1). We hypothesized that if M1 benefits from externally added noise in accordance to the SR phenomenon, neural responsiveness to transcranial magnetic stimulation (TMS) (i.e., reflecting the processing and/or transmission of the external stimulation) would be increased (McDonnell and Abbott 2009). Neural responsiveness was operationalized via the well-known concept of the resting motor threshold (RMT), which is defined as the stimulation intensity required to evoke motor evoked potentials (MEPs) of a given amplitude in at least 50% of trials, i.e., with a probability of 0.5. Accordingly, our hypothesis would be supported if RMTs were lower during tRNS application when compared to no stimulation because random noise increased the probability of evoking MEPs of sufficient size (**Figure 1**).

**Figure 1.**
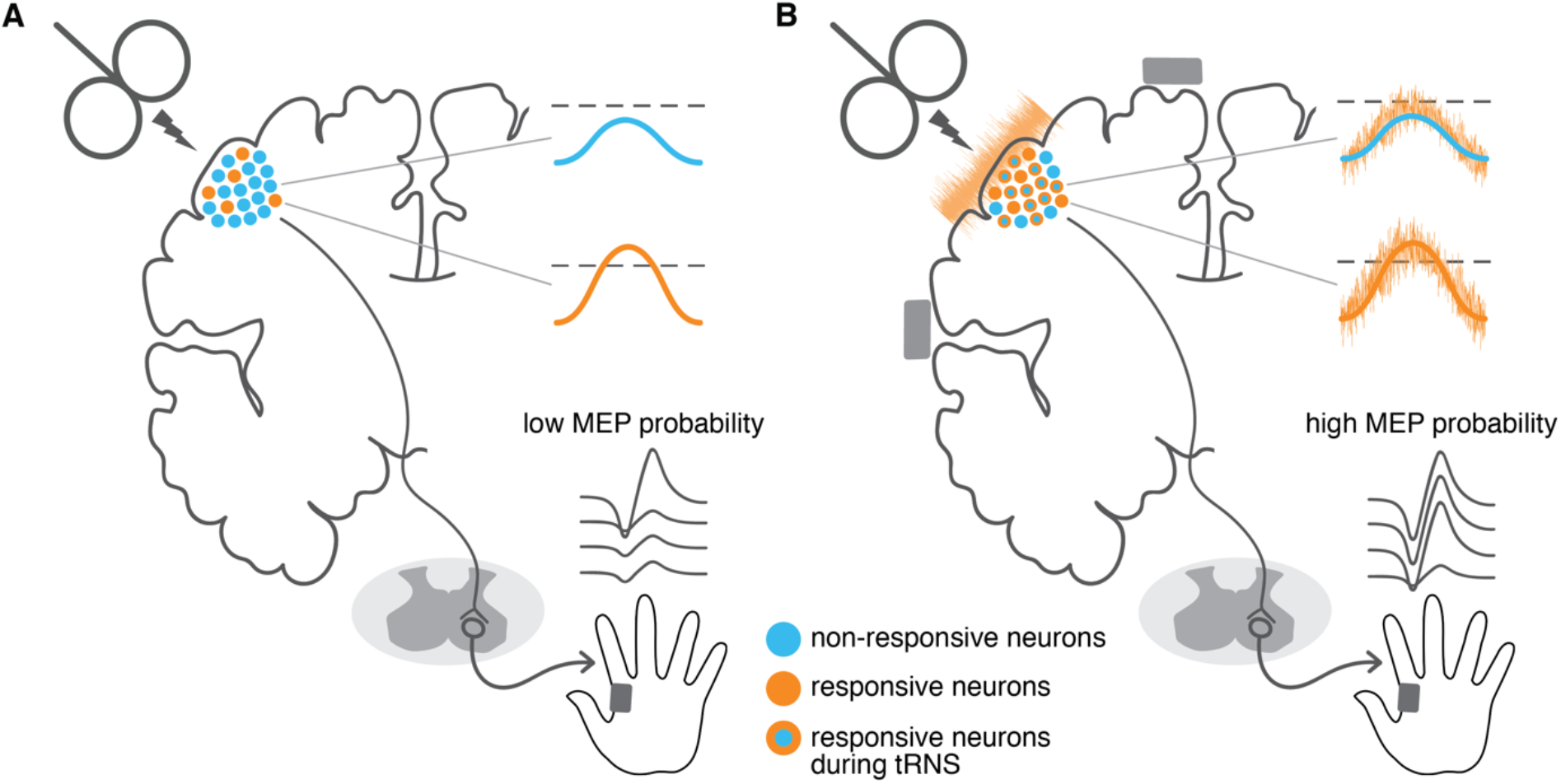
Conceptual representation of how transcranial random noise stimulation (tRNS) may enhance the neural signal and influence resting motor threshold (RMT). **A** Transcranial magnetic stimulation (TMS) pulse delivered with subthreshold intensity activates small population of neurons (orange circles) eliciting motor evoked potentials (MEPs) with a low probability. **B** When random noise is added to the primary motor cortex with tRNS (orange wave), neurons that did not respond before to TMS (blue circles) can cross the activation threshold (blue circles with orange outline). The enlarged portion of firing neurons results in higher probability of eliciting MEPs, which is reflected in a lower RMT.

## Materials and Methods

### Participants

Eighty-one healthy volunteers (46 females, 35 males, mean age = 25.4 ± 5.1; range, 18-46) took part in this study, which consisted of 4 experiments. A new group of participants was recruited for each experiment. All were right-handed and had no identified contraindications for participation according to established TMS exclusion criteria (Rossi et al., 2009; Wassermann, 1998). All provided written informed consent. Upon study conclusion, they were debriefed and financially compensated for their time and effort. None of the participants reported any major side effects resulting from the stimulation. All research procedures were approved by the Cantonal Ethics Committee Zurich (KEK-ZH-Nr. 2014-0242, KEK-ZH-Nr. 2014-0269 and BASEC Nr. 2018-01078) and were performed in accordance with the Helsinki Declaration of the World Medical Association (2013 WMA Declaration of Helsinki).

### General study design

To evaluate the acute influence of tRNS on the excitability of motor cortex we performed a series of four experiments in which we combined high frequency (100-500 Hz) tRNS with single-pulse TMS over left M1. TRNS was applied for only 3 seconds per trial, with jittering of the TMS pulse between 1.3 and 1.7s after tRNS onset (see **Figure 2** for an overview of the different experiments). Short tRNS duration was used based on the previously demonstrated acute behavioural effects of 2-5s tRNS (Groen and Wenderoth 2016; van der Groen et al. 2019) and reports showing that 4s tDCS results in an excitability change during stimulation without producing after-effects (Nitsche et al. 2003). In all experiments the TMS inter-trial-interval was set to 8 seconds with 20% temporal variability. During the measurement participants sat comfortably at rest and directed their gaze toward a fixation cross. Motor evoked potentials (MEPs) of the right first dorsal interosseous (FDI) muscle were used as a primary physiological read-out. Our main outcome parameter in all experiments was the probability of eliciting MEPs with an amplitude greater than or equal to 0.05 mV (i.e., response probability; p(MEP_0.05mV_)). All experiments consisted of conditions where tRNS was applied to M1 in random order (see below) and no noise control conditions, with the experimenter holding the TMS coil blinded regarding condition type. Since we used a very brief stimulation time (3 seconds only), fade in/out periods were not possible. Accordingly, some participants were able to distinguish the stimulation conditions (see Additional Control Analyses: Tactile Sensation). We accounted for this possible bias via various control analyses and by including one experiment where we introduced an active stimulation control condition (see Experiment 4).

**Figure 2.**
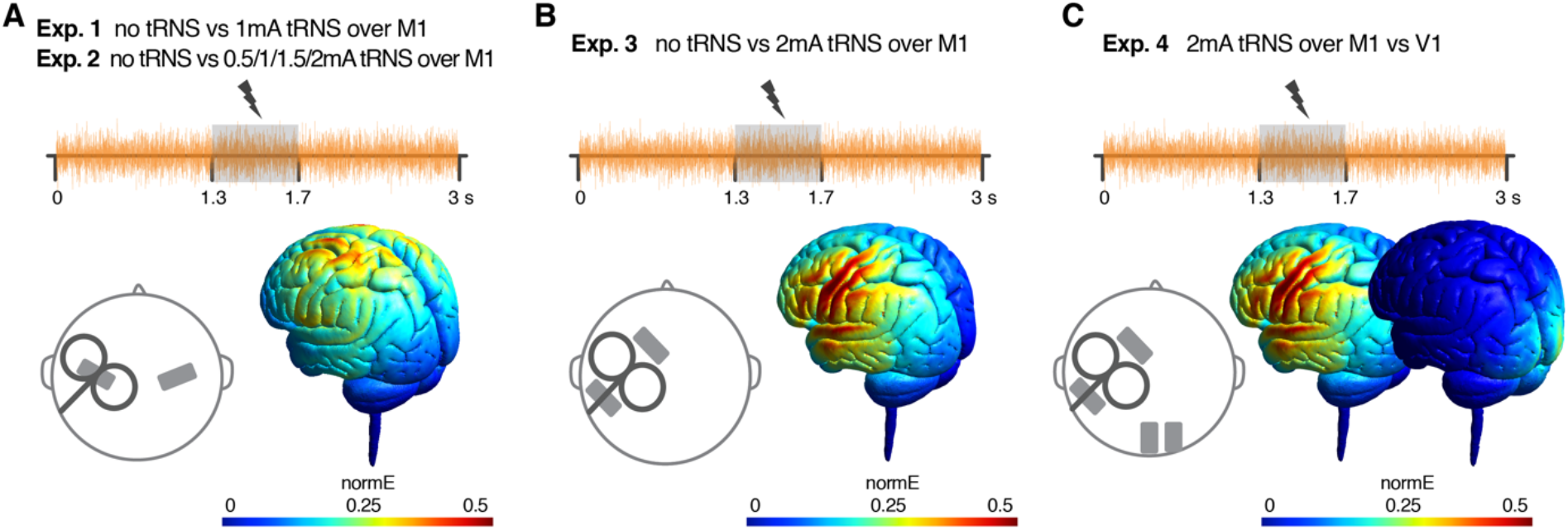
Stimulation protocol, electrode placement (grey rectangles) and electric field modelling for experiments 1-4. Transcranial random noise stimulation (tRNS, orange wave) was delivered online for 3 seconds over the primary motor cortex (M1) with transcranial magnetic stimulation (TMS) applied in the middle of the noise stimulation window. **A** In Exp.1 we delivered tRNS at 1mA (vs no tRNS) and probed motor evoked potentials (MEP) with TMS at resting motor threshold (RMT) intensity (jittered 1.3, 1.5 or 1.7 s after tRNS onset), to explore the noise influence on the probability of MEP. In Exp. 2 tRNS was delivered at variable intensity (0.5-2mA tRNS vs no tRNS) with subthreshold TMS (jittered, after 1.5 or 1.7 s) to test the noise dose-response effects. In both experiments, tRNS electrodes were placed over the left and right M1. Electric field modelling shows a 2mA noise intensity condition. **B** In Exp. 3 RMT_Fit_ was measured during the application of tRNS over M1 at 2mA (vs no tRNS) with a TMS threshold estimation approach (i.e., TMS applied in the range RMT ± 2%, jittered, 1.3 or 1.7 s after tRNS onset). TRNS electrodes were placed 7 cm anterior and posterior to the first dorsal interosseous (FDI) muscle hotspot (M1). **C** In Exp. 4 RMT_Fit_ was measured during the application of tRNS at 2mA over M1 (vs control stimulation site) with a TMS threshold estimation approach (i.e., TMS applied in the range RMT ± 2%, jittered, 1.3 or 1.7 s after tRNS onset). tRNS electrodes were placed anterior and posterior to the FDI muscle hotspot (M1 - main stimulation condition, left) and over the right occipital lobe (the control stimulation site, right).

### Transcranial Random Noise Stimulation

Random noise (100-500 Hz) was delivered through a battery-driven electrical stimulator (DC-Stimulator PLUS, NeuroConn GmbH, Ilmenau, Germany). Electroconductive gel was applied to the contact side of the rubber electrodes (5 × 7 cm) to reduce skin impedance. Depending on the experiment, the stimulation intensity varied between 0.5 - 2mA amplitude (peak-to-baseline), resulting in maximum current densities ranging from 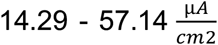, which is below the safety limits for transcranial electrical stimulation (Fertonani et al. 2015). The probability function of the stimulation followed a gaussian distribution with no offset. TRNS power, corresponding to the variance of the electrical noise intensities distribution (Thielscher and Saturnino 2019), was 0.73 mA^2^ in the 2mA condition. The impedance between the electrodes was monitored and kept below 15 kΩ (on average 8.6 ± 3.9 kΩ across experiments). TRNS waveforms were created in MATLAB (The MathWorks, Inc., Natick, USA), uploaded to Signal (Cambridge Electronic Design, version 2.13) and sent via a CED amplifier (CED Power 1401, Cambridge Electronic Design, Cambridge, UK) to the DC-stimulator which was operated in REMOTE mode. For all experiments we used electric field modelling to ensure optimal electrode placement (**Figure 2**). All simulations were run in SimNIBS 2.1 (Thielscher et al. 2015) using the average MNI brain template. The software enables finite-element modelling of electric field distribution of direct current stimulation without taking into account the temporal characteristics. Note, however, that the induced electric field was shown to be independent of the stimulation frequency (Vöröslakos et al. 2018). Since we were mainly interested in the peak of the induced electric field, we run the simulation for the maximum 2mA peak-to-baseline intensity of the stimulation.

TRNS and no noise control conditions were randomized throughout all of the experiments to prevent cumulative effects of tRNS and to minimize the effect of general changes in corticomotor excitability.

### Transcranial Magnetic Stimulation

Single-pulse monophasic TMS was delivered using a 70mm figure-of-eight coil connected to the Magstim 200 stimulator (Magstim, UK). For all subjects the coil was positioned over the hotspot of the FDI muscle. Depending on the tRNS electrode montage the coil was placed either on top of the electrode (Exp. 1 and 2) or on the scalp in between the electrodes (Exp. 3 and 4; see **Figure 2**). The hotspot was defined as the stimulation site where TMS delivery resulted in the most consistent and largest MEPs in the resting muscle. The coil was held tangential to the surface of the scalp with the handle pointing backward and laterally at 45° away from the nasion-inion mid-sagittal line, resulting in a posterior-anterior direction of current flow in the brain. Such a coil orientation is thought to be optimal for inducing the electric field perpendicular to the central sulcus resulting in the stimulation of M1 neurons (Mills et al. 1992; Rathelot and Strick 2009). The optimal coil location was marked with a semi-permanent marker on the head and registered using the neuronavigation software (Brainsight^®^ Frameless, Rogue Research Inc., Montreal, QC). The position of the participant’s head and TMS coil was constantly monitored in real-time with the Polaris Vicra^®^ Optical Tracking System (Northern Digital Inc., Waterloo, ON, Canada). This ensured that the centre of the coil was kept within 2 mm of the determined hotspot, and that the coil orientation was consistent throughout the experiment. For each participant we determined the resting motor threshold (RMT), defined as the lowest intensity to elicit MEPs with peak-to-peak amplitude greater than or equal to 0.05 mV in the relaxed muscle, in 5 out of 10 consecutive trials (Rossini, Barker, & Berardelli, 1994). Pooling data across all experiments, the mean RMT at baseline corresponded to 44 ± 9% of the maximum stimulator output (MSO).

### Electromyography

The muscle response was recorded by a surface electromyography (EMG) electrode (Bagnoli^™^ DE-2.1 EMG Sensors, Delsys, Inc.) placed over the right FDI muscle. Raw signals were amplified (sampling rate, 5 kHz), digitized with a CED micro 1401 AD converter and Signal software V2.13 (both Cambridge Electronic Design, Cambridge, UK), and stored on a personal computer for off-line analysis. Timing of the TMS delivery, remote control of the tRNS stimulator and EMG data recording were synchronized via the CED. Muscular relaxation was constantly monitored through visual feedback of EMG activity and participants were instructed to relax their muscles if necessary.

### Data processing and analysis

The EMG data was band-pass filtered (30-800 Hz, notch filter = 50 Hz). Filtering was applied separately for the pre-TMS background EMG (bgEMG) and post-TMS period containing peak-to-peak MEP amplitude in order to avoid “smearing” the MEP into bgEMG data. Peak-to-peak amplitude was defined as the peak-to-peak amplitude between 15 to 60 ms after the TMS pulse. Next, we excluded trials in which the unwanted background muscle activation could influence the measured MEP amplitude. Trials with root mean square bgEMG above 0.01 mV were removed from further analyses to control for unwanted bgEMG activity (Hess et al., 1986; Devanne et al., 1997). For the remaining trials, the mean and standard deviation of the background EMG was calculated for each participant. Trials with bgEMG > mean ± 2.5 standard deviations were also excluded. Based on these criteria 96.7% of all data were included for further analysis (96.7% in Exp.1, 97.7% in Exp. 2, 96.2% in Exp. 3 and 97.6% in Exp. 4).

In all experiments our main outcome parameter was the probability of eliciting MEPs with an amplitude greater than or equal to 0.05 mV (for each condition 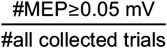; p(MEP_0.05mV_)). We decided to investigate RMT modulation since it allows us to assess the individual membrane excitability of the corticospinal tract neurons (Kobayashi and Pascual-Leone 2003; Hallett 2007; Rossi et al. 2009). RMT serves as one of the most robust TMS measurements (Kobayashi and Pascual-Leone 2003; Nitsche et al. 2005; Hallett 2007; Livingston and Ingersoll 2008; Rossi et al. 2009; Ngomo et al. 2012; Hinder et al. 2014; Schambra et al. 2015; Davila-Pérez et al. 2018; Dissanayaka et al. 2018; Jannati et al. 2019). For all experiments we ran additional control analyses by re-calculating MEP probabilities for other amplitude criteria, i.e., MEP amplitudes larger than or equal to 0.03, 0.04, 0.06 and 0.07 mV to ensure that the observed effects are not purely driven by the definition of the RMT. Additionally, we performed a supplementary analysis to investigate the potential influence of the noise stimulation on MEP amplitude.

### Statistical analysis

Statistical analyses were performed with repeated measures ANOVAs (rmANOVA) in IBM® SPSS Statistics Version 23.0 (IBM Corp., Armonk, NY, USA) unless otherwise stated. All data were tested for normal distribution using Shapiro-Wilks test of normality. Sphericity was tested using Mauchly’s sphericity test. If sphericity was violated, Greenhouse-Geisser correction was applied. The threshold for statistical significance was set at α = 0.05. All post hoc tests were corrected for multiple comparisons using Bonferroni correction. Effect sizes are reported for each experiment in the form of Partial Eta Squared (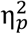 small 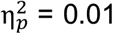, medium 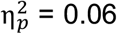, large 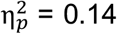 Lakens, 2013) or Cohen’s d (d_z_; small d_z_ = 0.2, medium d_z_ = 0.5, large d_z_ = 0.8; Lakens, 2013). Variance is reported as standard deviation (SD) in the main text and as standard error (SE) in the figures. Potentially confounding variables (i.e., assessed tactile sensation and bgEMG) were added as covariates whenever applicable (see **Additional Control Analyses** in the results section).

### Additional information for specific experiments

#### Experiment 1 – Testing effects of 1mA tRNS

In the first experiment, we tested whether tRNS induces an increase in MEP probability when TMS was applied to M1 at RMT intensity. Sixteen right-handed participants (self-report) took part in the experiment (9 females, 7 males, mean age = 24.7 ± 5, range: 19-35). TRNS electrodes were placed over (i) the hotspot of the right FDI as determined by single-pulse TMS and (ii) the contralateral, right M1 (**Figure 2A**). This bilateral montage was chosen based on modelling results which indicated similar current densities but less current spread for the M1-M1 montage (used here) compared to the M1-supraorbital cortex montage as used in previous studies (Terney et al. 2008; Chaieb et al. 2011). TRNS intensity was set to 1mA peak-to-baseline amplitude and applied in each trial for 3 seconds. TMS pulses were delivered over left M1 at RMT intensity (mean RMT 49 ± 13 %MSO, range: 36-84%), starting 1.3, 1.5 or 1.7 s after tRNS onset. Testing was split into 4 blocks (60 trials in each block, 240 trials total). We measured MEPs in the FDI muscle either during the 1mA tRNS condition (180 trials total) or the no tRNS condition (control, 60 trials total). We compared MEP probability between the 1mA tRNS vs no tRNS conditions using a paired t-test. Even though we adjusted the TMS intensity to the individuals’ RMT prior to the main experiment, closer inspection of the data revealed that RMTs could drift during the experiment (Karabanov et al. 2015) so that some participants were stimulated at sub-threshold TMS intensities (i.e., MEP_0.05mV_ probability < 0.5 in the no tRNS condition) while others were stimulated at supra-threshold intensities (MEP_0.05mV_ probability > 0.5 in the no tRNS condition). Therefore, we evaluated whether tRNS-induced changes in MEP probability were associated with MEP probability in the no tRNS condition using Pearson correlation. The tRNS-induced modulation was calculated as a ratio between p(MEP_0.05mV_) in the 1mA tRNS condition and p(MEP_0.05mV_) in the no tRNS condition. Thus, we correlated *x* with 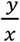, where:

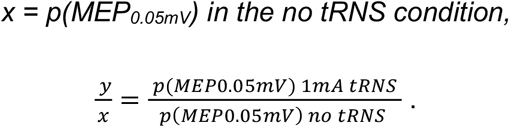

Given that correlating a fractional increase from x with x itself can be problematic as it might lead to spurious correlations (Pearson 1897; Tu 2016), we applied a statistical correction method suggested by Tu (2016; Eq. (4-6)). This method compares the above correlation r_x,y/x_ to the expected “null correlation” (r_null_) which is revealed by

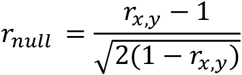

The observed correlation r_x,y/x_ is then compared to r_null_ using a z-test of the following form:

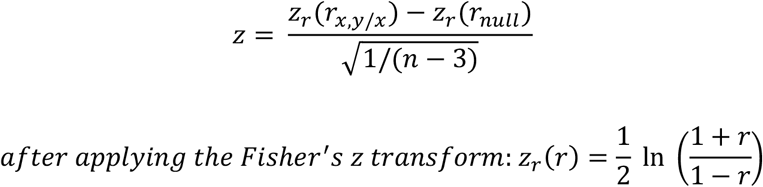

This allowed us to assess if there is a statistically significant association between p(MEP_0.05mV_) in the no tRNS condition and tRNS-induced modulation.

#### Experiment 2 – Testing dose-response effects

The second experiment aimed to determine whether there is an optimal tRNS intensity to modulate MEP probability in M1. Twenty-three right-handed (Edinburgh Handedness Inventory mean laterality quotient [LQ] = 78.3 ± 13.8; Oldfield, 1971) participants were recruited. We had to exclude one participant because of technical problems with data acquisition. Twenty-two participants (13 females, 9 males, mean age = 25.4 ± 5.4, range: 20-46) were included in the subsequent analysis. At the beginning of the session we measured the RMT of each participant (mean RMT = 43.4 ± 7.5 %MSO, range: 31-54%). Based on the results of experiment 1 and behavioural effects of tRNS (Groen and Wenderoth 2016; van der Groen et al. 2018) we decided to probe MEP with subthreshold TMS, as the response towards stimuli presented below the threshold are postulated to be particularly susceptible for SR effects (Gingl et al. 1995; Moss et al. 2004). TMS over left M1 was always applied with an intensity slightly below RMT, which was operationally defined as the intensity evoking MEPs with an amplitude of at least 0.05 mV in 3 out of 10 trials (p(MEP_0.05mV_) = 0.3; mean subthreshold intensity: 42 ± 7.3 %MSO, range: 30-53%). The TMS pulse was randomly applied either 1.5 or 1.7 s after tRNS onset to avoid anticipation of the pulse (**Figure 2A**). The tRNS electrodes were placed as in experiment 1 (**Figure 2A**). We varied tRNS intensity (0.5, 1, 1.5 or 2mA vs no tRNS control condition) to examine whether there is a dose effect on the tRNS induced enhancement of MEP probability. We recorded 10 trials per tRNS condition in randomized order in 3 blocks. Throughout the experiment we monitored MEP probability in the no tRNS control condition and, if required, we adjusted the TMS intensity between blocks to ensure subthreshold level stimulation (p(MEP_0.05mV_) = 0.3, **Figure 9**). This procedure was chosen because it has been shown that state-dependent changes of RMT can occur in absence of overt activity or task involvement and need to be considered in order to keep TMS intensity comparable throughout the experiment (Karabanov et al. 2015). Therefore, we determined MEP probability for the no tRNS condition after each completed block. Since we targeted MEP probability of 0.3, TMS intensity was reduced by 1 %MSO when MEP probability was ≥ 0.5 or increased when MEP probability was ≤ 0.1. Note that adjusting TMS intensities only between blocks ensured subthreshold TMS throughout the course of the experiment without confounding the comparison between the different tRNS conditions that were sampled equally within each block. 30 MEPs were collected per condition resulting in a total number of 150 TMS pulses. For nineteen participants we collapsed MEPs across the three blocks to estimate the MEP probability for each of the noise conditions. For three participants, post-hoc analysis revealed that they met the criteria for subthreshold TMS during the no tRNS control condition only in two blocks, i.e., MEP probabilities were estimated from 20 MEPs per condition. MEP probabilities were subjected to a rmANOVA with the within-subject factor *tRNS intensity* (no tRNS, 0.5, 1, 1.5 and 2mA tRNS).

**Figure 9.**
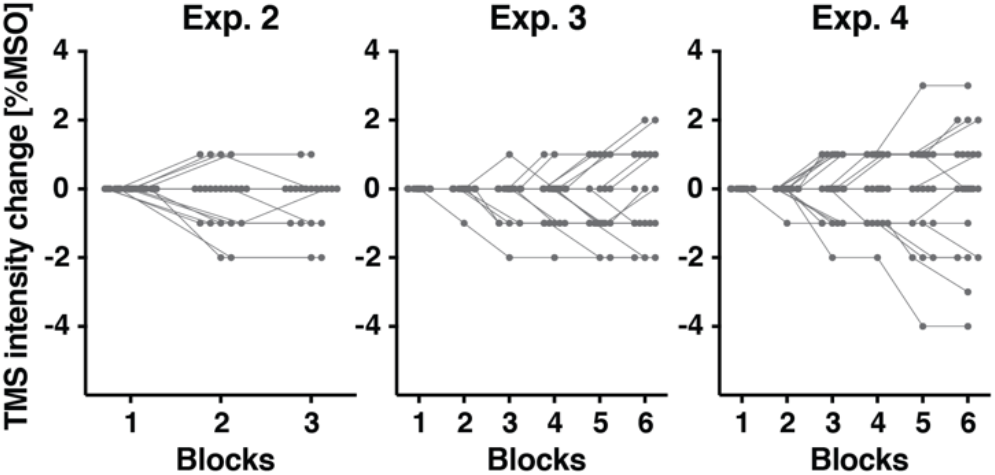
Changes in the adjusted transcranial magnetic stimulation (TMS) intensity between measurement blocks in experiments 2-4. Grey dots represent TMS intensity change for each individual between the different blocks of\ experiments 2-4.

Before starting the main experiment, all experimental tRNS intensities (0.5-2mA) were presented to the participant (for 20 s in a randomized order) and subjectively assessed on a scale from 0 (no sensation) to 10 (strong pain) to make sure that the stimulation did not cause any unpleasant sensations.

#### Experiment 3 – Threshold estimation

The third experiment was a conceptual replication of experiment 2, but this time we used a modified electrode montage and tested whether tRNS applied online can influence RMT. The required sample size was estimated based on Experiment 2 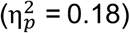 using a power analysis (G* Power version 3.1; Faul, Erdfelder, Lang, & Buchner, 2007). It revealed that twenty participants should be included to detect an effect of tRNS on MEP probability with a 2×5 rmANOVA, alpha = 0.05 and 85% power. Twenty-three right-handed (mean LQ = 88.7 ± 16.5; Oldfield, 1971) individuals were recruited in experiment 3 to account for potential dropouts, with three removed from further analyses (see below).

RMT was defined at the beginning of the session (mean RMT = 41.1 ± 7.4 %MSO, range: 28-55%). We then applied TMS intensities centred around the individual RMT level during the main experiment (namely: RMT-2%, RMT-1%, RMT, RMT+1%, RMT+2%). Based on the results of experiment 2, we tested only tRNS intensity of 2mA versus no tRNS. This time we used a modified electrode montage as described by Rawji et al. (2018), whereby electrodes were positioned 7 cm anterior and posterior to the FDI muscle hotspot along the coil axis (45° away from the nasion-inion mid-sagittal line). As described by the authors, in this arrangement current oscillates perpendicular to the central sulcus, which has been hypothesized to be more efficient in modulating corticospinal excitability (Rawji et al. 2018). The electric field modelling showed that this arrangement provides more focal electrical stimulation and induces a slightly higher electric field over M1 (**Figure 2B**). Additionally, this montage enables positioning the TMS coil directly on the scalp and avoids delivering TMS pulses through the electrode (see **Figure 2**). Different electrode montages can change the directionality of the electric field which has been shown to strongly influence the effect of tDCS (Nitsche et al. 2008). However, noise benefits induced by tRNS should be polarity independent (Pirulli et al. 2016). Here we test whether the increased M1 responsiveness caused by applying tRNS generalize across electrode montages targeting M1. We tested 10 different experimental conditions (no tRNS versus 2mA tRNS x 5 TMS intensities) in total. The experiment consisted of 6 blocks, with 4 trials per condition presented within each block in a randomized order (24 MEPs in each of the 10 conditions, 240 TMS pulses total). TMS was randomly applied 1.3 or 1.7 s after tRNS onset within each trial (**Figure 2B**). We monitored the MEP probability in the control condition (no tRNS with TMS intensity targeted at RMT) and TMS intensity was adjusted between blocks by 1 %MSO: it was (i) decreased if the probability of MEPs in the no tRNS condition was ≥ 0.75 or (ii) increased if the probability of MEPs was ≤ 0.25, to stay as close as possible to the target RMT level (p(MEP_0.05mV_) = 0.5; **Figure 9**).

Post-hoc inspection of the data revealed that two participants showed response probabilities consistently above RMT. Thus, they did not fulfil our criteria of probing MEP with TMS intensities centred around the RMT and were excluded from subsequent analyses. Additionally, one outlier was excluded during data analysis due to a tRNS-induced decrease of RMT > 2 SD of the group mean. Data from the final sample of twenty participants (10 females, 10 males, mean age = 27.5 ± 6, range: 18-42) was entered into a 2×5 rmANOVA with the within-subject factors *tRNS* (no tRNS versus 2mA tRNS) and *TMS intensity* (RMT-2%, RMT-1%, RMT, RMT+1%, RMT+2%). Additionally, we performed a threshold estimation analysis to determine whether tRNS influenced RMT. To do so, for each participant we calculated the response probability for each of the 5 TMS intensities when either 2mA tRNS or no tRNS was applied. Next, we fitted separate linear models (*y = ax + b*, with y denoting *p(MEP*_*0*.*05mV*_) and x the stimulation intensity) to each of the datasets and determined *RMT*_*Fit*_ *= (0*.*5-b)/a*, i.e., the intensity which would be used to evoke a sufficiently large MEP with a probability of 0.5 (see **Figure 3**). Note that this method provides a more accurate estimation of RMT than manually adjusting TMS intensity until p(MEP_0.05mV_) = 0.5 is reached, partly because the model is informed by more data.

**Figure 3.**
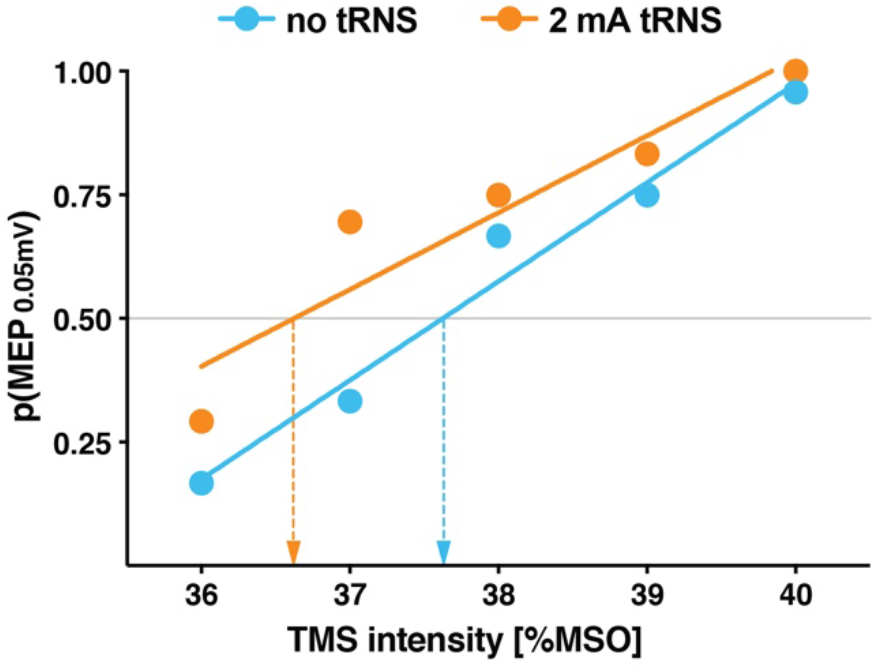
Representative data of an individual participant to exemplify the threshold estimation procedure. The probability of eliciting motor evoked potential (p(MEP_0.05mV_)) was determined for five intensities ranging from RMT-2% to RMT+2% (RMT corresponds here to 38 %MSO). These values were obtained for the no tRNS control condition (blue symbols) as well as during 2mA tRNS over primary motor cortex (orange symbols). A linear model (y = ax+b) was fitted to the data of each condition (represented by the solid lines) and we determined which stimulation intensity would yield p(MEP_0.05mV_) = 0.5 via the following formula RMT_Fit_ = (0.5-b)/a. The figure symbolizes this procedure by showing that an intensity of RMT_Fit_ = 37.63% was determined for the no tRNS condition (blue arrow), while a slightly smaller RMT_Fit_ = 36.62% was determined for the 2mA tRNS condition (orange arrow).

Before the start of the main experiment, participants were familiarized with tRNS and we assessed the detectability of potential sensations. The detection task consisted of 20 trials. Participants received either tRNS (2mA over M1, with and without TMS on half of the trials) or no tRNS (with and without TMS). Their task on each trial was to indicate (after an auditory cue) if they felt something underneath the tRNS electrodes (ignoring TMS pulses) by pressing the appropriate button on a keyboard.

#### Experiment 4 – Threshold estimation with active control condition

The final experiment aimed to replicate the results obtained in experiment 3, but this time we compared tRNS applied over M1 to tRNS applied over right primary visual cortex (V1) to control for unspecific stimulation effects. The sample size estimation was based on the effect size of 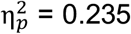 which was obtained by averaging the effect sizes of the tRNS main effect in experiment 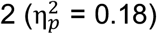 and 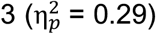 to get a robust estimation across experiments. We further assumed that sphericity might be violated (ε = 0.55 as in Exp. 3) and set alpha = 0.05 and power = 85%. This revealed a sample size of twenty-two participants. Based on this analysis we recruited twenty-nine individuals to account for potential dropouts (see below). Experiment 4 used the same stimulation parameters, electrode montage and general procedures as experiment 3, but this time the no tRNS control condition was replaced with 2mA tRNS over the right occipital lobe. The control site was selected to evoke similar skin sensations as the main experimental condition (i.e., applying 2mA tRNS over M1), but to deliver stimulation that does not interfere with neural processing related to the measured MEPs (see the electrical field modelling, **Figure 2C**). The first electrode was placed over the inion and the second electrode was placed to the right (7 cm between the centres of the electrodes, **Figure 2C**). TRNS was delivered with a separate battery-driven remote-controlled electrical stimulator (DC-Stimulator PLUS, NeuroConn GmbH, Ilmenau, Germany), with the same high frequency (100-500 Hz) tRNS waveform of 2mA intensity (peak-to-baseline amplitude with a 0mA offset) created in MATLAB (The MathWorks, Inc., Natick, USA).

We applied tRNS either over the left M1 or the control stimulation site and probed the muscle response with single-pulse TMS using 5 intensities around threshold level (RMT-2%, RMT-1%, RMT, RMT+1%, RMT+2%). As in experiment 3, the TMS intensity was adjusted between blocks if necessary, based on the data obtained during the control stimulation over right V1 (**Figure 9**).

From the original sample of twenty-nine right-handed (mean LQ = 85 ± 20.3; Oldfield, 1971) participants, six were excluded for various reasons. One could not complete the session due to technical problems. Three participants revealed response probabilities consistently above RMT and one substantially below RMT during the experiment. Additionally, one outlier was removed post-hoc because tRNS caused the RMT to decrease by more than 2 SD of the group mean. This resulted in the final sample of twenty-three participants (14 females, 9 males, mean age = 24.3 ± 3.9, range: 19-34; mean RMT = 44.1 ± 7.2 %MSO, range: 33-59%). Similar to the previous experiment, we used the threshold estimation analysis (RMT_Fit_) and a 2×5 rmANOVA with the within-subject factors of *tRNS* (tRNS over M1 vs V1) and *TMS intensity* (RMT-2%, RMT-1%, RMT, RMT+1%, RMT+2%) for the statistical analysis of MEP probability. As in experiment 3, we assessed the detectability of skin sensation due to electrical stimulation via a detection task, which was performed before and after the main experiment. Detection task was similar to experiment 3, but included three stimulation conditions: no tRNS, 2mA tRNS over M1 or 2mA tRNS over right V1.

## Results

### Exp. 1 – TRNS induced increase in MEP probability for subthreshold TMS

In the first experiment we investigated whether tRNS modulates the probability of eliciting MEPs. We measured MEPs during 1mA tRNS versus no tRNS and calculated the probability of evoking MEPs with an amplitude ≥ 0.05 mV (p(MEP_0.05mV_) 1mA tRNS and p(MEP_0.05mV_) no tRNS, respectively). We did not observe a significant difference between the overall MEP probability in the noise vs control condition (t_(15)_ = 0.31, p = 0.77, mean difference [MD] = 0.007 ± 0.09).

Even though we aimed for stimulating each participant with a TMS intensity corresponding to RMT as determined prior to the main experiment, post-hoc analysis of the no tRNS condition revealed that some participants were stimulated with subthreshold TMS intensities (i.e., p(MEP_0.05mV_) no tRNS < 0.5) while others were stimulated with suprathreshold TMS intensities (i.e., p(MEP_0.05mV_) no tRNS > 0.5). Therefore, we calculated a Pearson correlation to test whether a potential increase in p(MEP_0.05mV_) in the 1mA tRNS condition (i.e., indicating a noise benefit) depended on the MEP probability in the no tRNS condition (**Figure 4**). We found a clear negative correlation r_x,y/x_ = −0.68, which was highly significant when compared to the null model r_null_= 0.15 (z = −3.56, p < 0.001, see methods for details). This finding suggests that random noise stimulation modulates M1’s response probability most strongly when TMS stimuli are delivered with an intensity slightly below the individual RMT (i.e., p(MEP_0.05mV_) no tRNS < 0.5), a result that is consistent with van der Groen et al. (2016) who showed that tRNS enhances detection performance for subthreshold stimuli.

**Figure 4.**
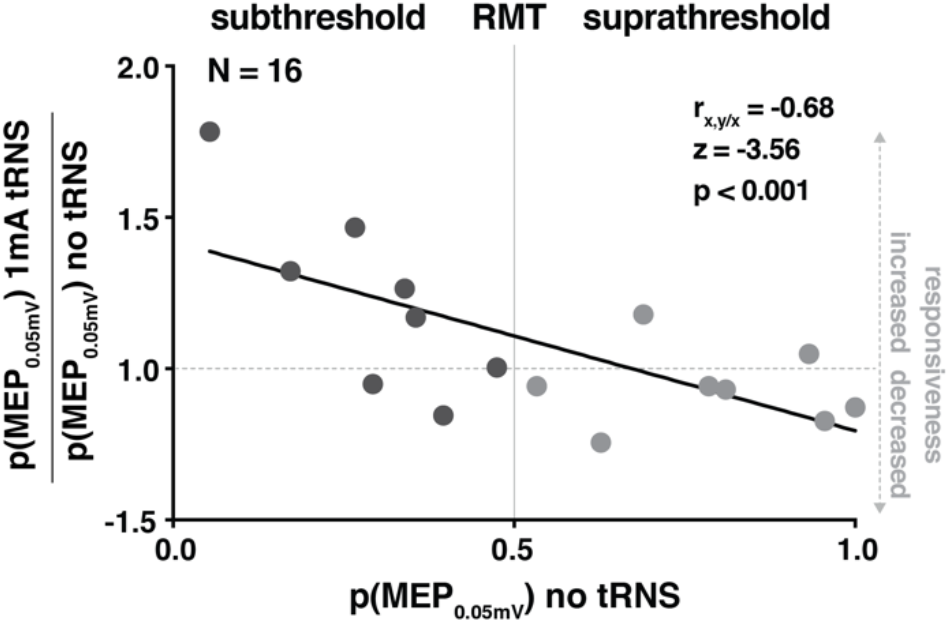
Transcranial random noise stimulation (tRNS) induced an increase in motor evoked potential (MEP) probability relative to the probability of eliciting MEPs in the no tRNS control condition. The Y axis represents MEP probability in the noise condition normalized to the individual MEP probability in the no tRNS condition, i.e., *p(MEP*_*0*.*05mV*_*) 1mA tRNS / p(MEP*_*0*.*05mV*_*) no tRNS*. Participants that were stimulated with subthreshold TMS intensities (i.e., *p(MEP*_*0*.*05mV*_*) no tRNS < 0*.*5*, dark grey symbols on the left) benefited more from 1mA tRNS than participants that were stimulated at suprathreshold TMS intensities (i.e., *p(MEP*_*0*.*05mV*_*) no tRNS > 0*.*5*, light grey symbols on the right). Grey dots indicate single subject data. Statistics for r_x,y/x_ are reported relative to the null correlation (r_null_), see methods for details.

### Exp. 2 - increase in MEP probability for higher tRNS intensity over M1

Next, we aimed to examine if there is an optimal tRNS intensity to enhance the probability of evoking MEPs. Therefore, we applied random noise stimulation over both M1s at intensities ranging from 0.5, 1, 1.5 to 2mA vs no tRNS control condition (**Figure 2A**). Based on our previous finding, we ensured that M1 was probed with TMS at sub-RMT intensities. Accordingly, p(MEP_0.05mV_) = 0.26 ± 0.06 for the no tRNS condition but gradually increased for higher tRNS intensities as indicated by a significant main effect of tRNS intensity (F_(4, 84)_ = 4.57, p = 0.002, 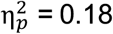, **Figure 5**). Post-hoc comparisons revealed that the 2mA stimulation was most effective in boosting MEP probability, which differed significantly from the no tRNS control condition (p = 0.04, MD = 0.082 ± 0.1) and 0.5mA stimulation (p = 0.03, MD = 0.079 ± 0.1). This indicates that the probability of inducing MEPs scales with increasing tRNS intensities.

**Figure 5.**
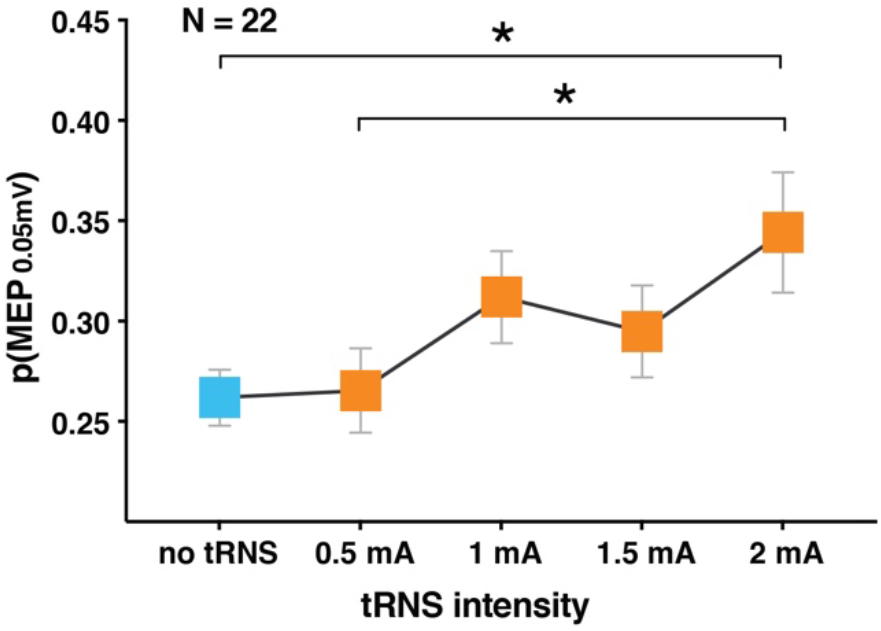
Probability of eliciting motor evoked potentials (p(MEP_0.05mV_)) at different transcranial random noise stimulation (tRNS) intensities, revealing a significant increase in the probability of evoking a MEP during 2mA tRNS. Error bars indicate SE. * indicates p < 0.05.

### Exp. 3 - TRNS over M1 induced decrease in RMT

Next, we performed a conceptual replication of the previous experiment and applied 2mA tRNS versus no tRNS (control condition) but changed the electrode placement (**Figure 2B**) to probe whether acute noise benefits on responsiveness generalize across different electrode montages targeting M1. Even though specific electrode positions can possibly affect the directionality of the electrical field to target neurons and result in divergent effects regarding polarity and potentially also the effective amount of the induced current, we again observed tRNS-induced enhancement in cortical responsiveness. We found a general increase in MEP probability when 2mA tRNS was applied over M1 (main effect of tRNS, 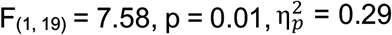, **Figure 6A**). The interaction between tRNS and TMS intensity was not significant (F_(4, 76)_ = 2, p = 0.11). Next, we calculated RMT_Fit_ as an additional outcome parameter. Note that RMT is tightly related to MEP probability since it is defined as the intensity which evokes sufficiently large MEPs with p(MEP_0.05mV_) = 0.5. We estimated RMT_Fit_ for each condition in each individual and found that RMT_Fit_ was significantly lower when 2mA tRNS versus no tRNS was applied (t_(19)_ = 2.3, p = 0.03, MD = 0.37 ± 0.73, d_z_ = 0.51; **Figure 6B**). This effect was generally small (≤ 2.1 %MSO), but relatively consistent across individuals as 14 out of 20 participants exhibited a slight decrease in RMT (**Figure 6C**). The results confirm that tRNS influences cortical responsiveness, which was reflected by a lower threshold at rest.

**Figure 6.**
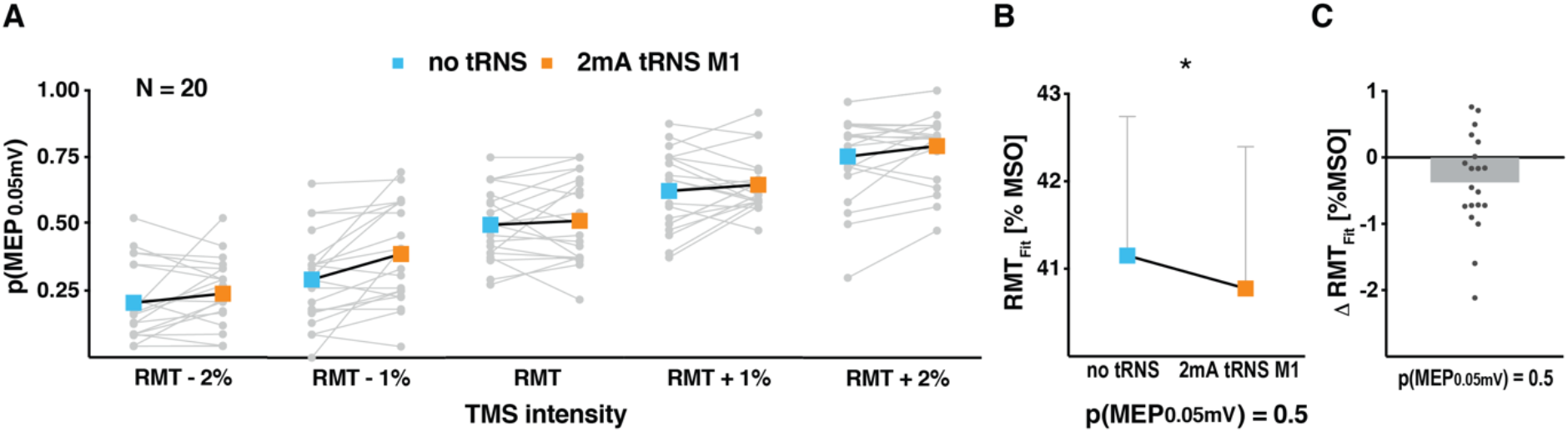
Increase in motor evoked potential probability (p(MEP_0.05mV_)) and decrease in the fitted resting motor threshold (RMT_Fit_) during transcranial random noise stimulation (tRNS) over primary motor cortex (M1) in comparison to the no tRNS control condition. **A** Single-subject data and average change in MEP probability in the no tRNS control and tRNS conditions at different levels of transcranial magnetic stimulation (TMS). **B** Decrease in RMT_Fit_ during tRNS over M1 in comparison to the no tRNS control condition. RMT_Fit_ refers to the TMS intensity needed to obtain a 0.5 MEP probability level in both conditions. Individual RMT_Fit_ values were assessed in the threshold estimation analysis based on responses from A. **C** Modulation of RMT_Fit_: individual differences between RMT_Fit_ in the 2mA tRNS and no tRNS condition (from B). Error bars indicate SE, grey dots indicate single subject data, grey bar indicates group mean, * indicates p < 0.05.

### Exp. 4 - decrease in RMT is specific for M1 stimulation

In experiment 4 we enrolled a new cohort of participants to control for the potentially unspecific effects of tRNS (e.g. arousal or tactile stimulation; Fertonani et al., 2015) by comparing 2mA tRNS over M1 to 2mA tRNS over right V1, with the latter serving as a control area that is unlikely to influence RMT. Our results confirmed the principal finding from the previous experiment, revealing that the probability of evoking MEPs is higher when stimulating M1 compared to the control site (main effect of tRNS, 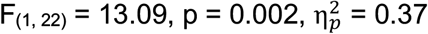, but no significant tRNS x TMS intensity interaction, F_(4, 88)_ = 0.42, p = 0.8; **Figure 7A**). We found that RMT_Fit_ was lower when 2mA tRNS was delivered over left M1 compared to right V1 (t_(22)_ = 4.5, p < 0.001, MD = 0.67 ± 0.71, d_z_ = 0.94; **Figure 7B**). Even though the absolute decrease was small (≤ 2.3 %MSO), 20 out of 23 participants exhibited a slight reduction in RMT_Fit_. Our results demonstrate that 2mA tRNS modulates the responsiveness of cortical motor circuits, as indicated by the decrease in individual motor threshold, an effect that is specific for the stimulation of M1..

**Figure 7.**
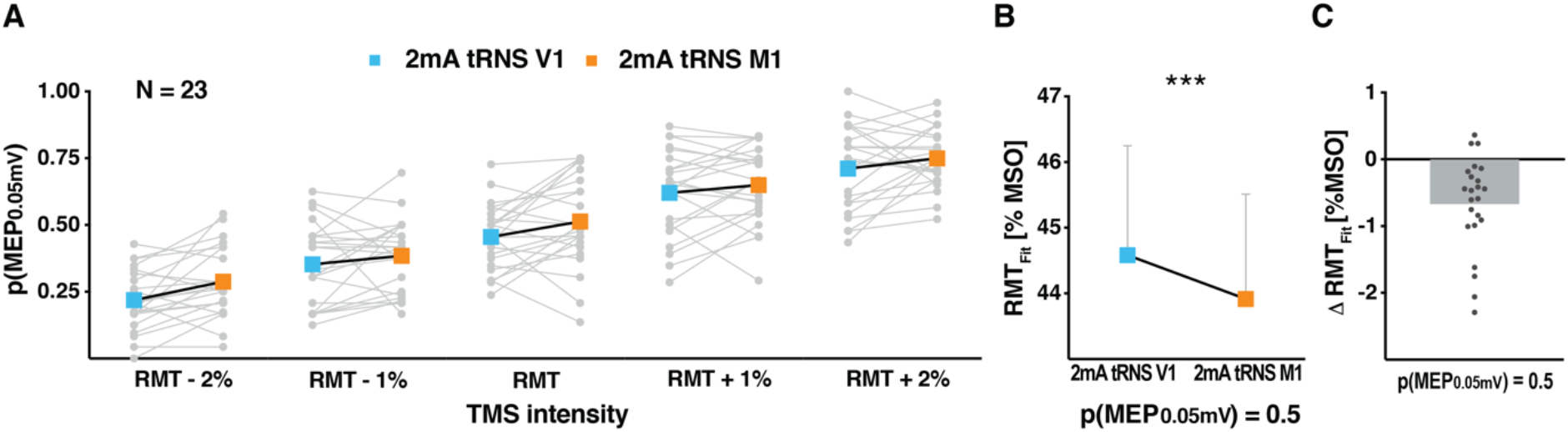
Increase in motor evoked potential probability (p(MEP_0.05mV_)) and decrease in the fitted resting motor threshold (RMT_Fit_) during noise stimulation over primary motor cortex (M1) in comparison to the control stimulation condition. **A** Single-subject data and average changes in MEP probability during noise stimulation over M1 and the control site at different levels of transcranial magnetic stimulation (TMS). **B** Decrease in RMT_Fit_ during tRNS application over M1 in comparison to the control site. RMT_Fit_ refers to the estimated TMS intensity needed to obtain a 0.5 MEP probability level in both conditions. Individual RMT_Fit_ values were assessed in the threshold estimation analysis based on responses from A. **C** Modulation of RMT_Fit_: individual differences between RMT_Fit_ during 2mA tRNS over M1 and the control site (from B). Error bars indicate SE, grey dots indicate single subject data, grey bar indicates group mean, * * * indicates p < 0.001

### Additional control analyses

#### MEP Amplitude

We analysed whether tRNS over M1 at 0.5mA up to 2mA versus the control condition (i.e., no tRNS or 2mA tRNS over right V1) influenced average MEP amplitude. Over all experiments this effect did not reach significance (all p ≥ 0.23), except for experiment 3 where we found a significant increase in MEP amplitude in the 2mA tRNS vs no tRNS condition (main effect of tRNS, 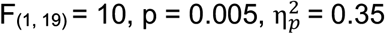; **Figure 8**).

**Figure 8.**
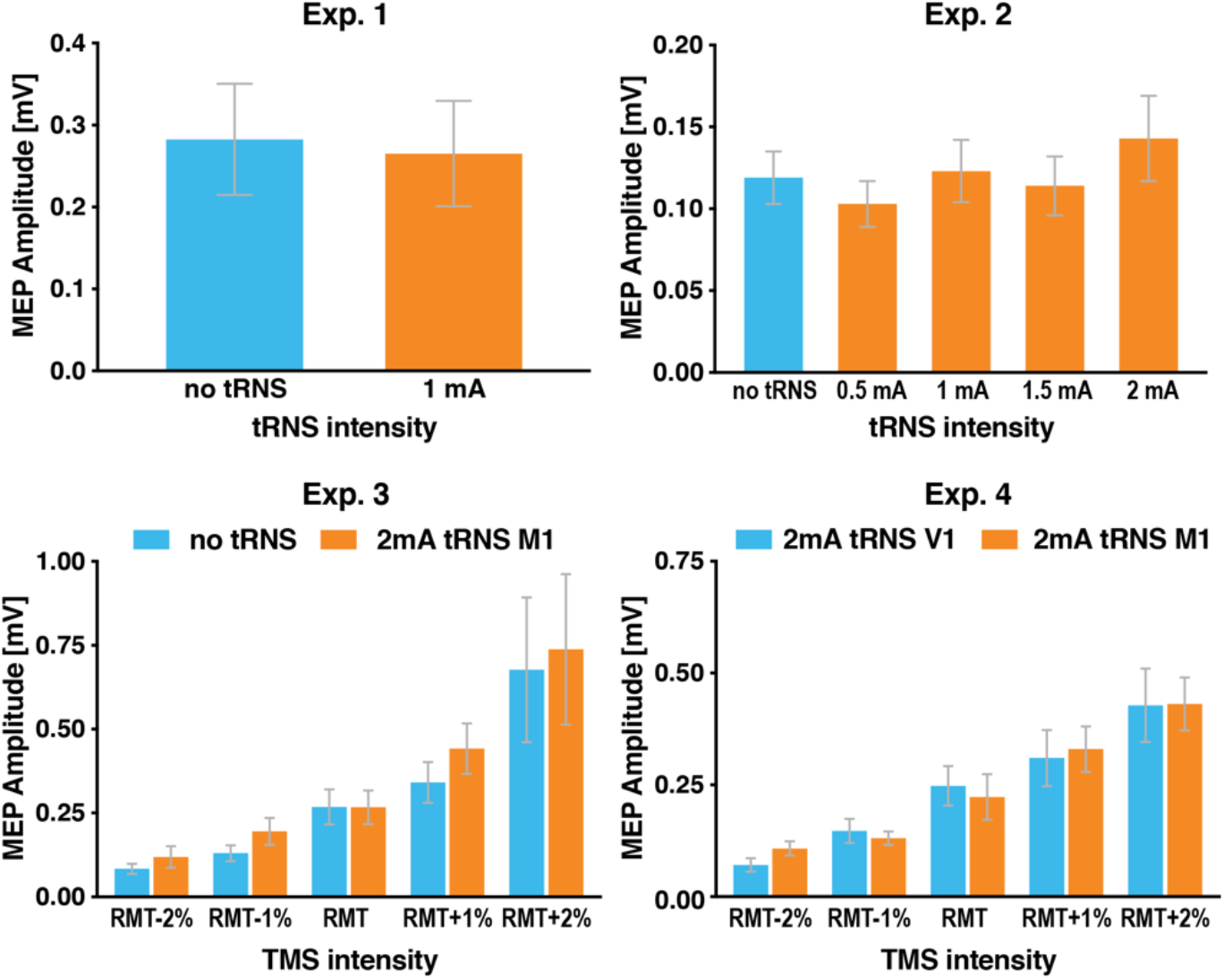
Average motor evoked potential (MEP) amplitude elicited by single-pulse transcranial magnetic stimulation (TMS) in the experimental and control conditions. tRNS modulated MEP amplitude to a minor extent resulting in significant effects only in Exp. 3, but not in Exp. 1, 2 and 4. Error bars indicate SE.

#### Tactile sensation

In experiments 2-4 we performed assessments of the possible tactile effects caused by electrical stimulation. We found that most of our participants could distinguish between no tRNS and tRNS conditions. In experiment 2 we recorded the subjective assessment (on a 0-10 scale) of the tactile sensation evoked by tRNS (mean for 0.5mA = 0.3 ± 0.7; 1mA = 0.6 ± 1.4; 1.5mA = 1.2 ± 1.8; 2mA = 1.9 ± 1.9). In experiment 3 we measured the detectability of potential sensations due to 2mA tRNS via a detection task (mean accuracy = 89% ± 18). We extended the detectability estimation in experiment 4, by repeating the task before (Pre) and after (Post) the experiment (mean accuracy Pre = 85% ± 17; mean accuracy Post = 77% ± 18, resulting in general average accuracy of 80% ± 16). Additionally, we distinguished hit rates (HRs) for correct detection of M1 (mean = 0.71 ± 0.4) versus V1 stimulation (mean = 0.74 ± 0.3), showing that sensation detectability did not significantly differ between stimulated areas (t_(22)_ = −0.27, p = 0.79). In order to test whether our TMS results might have been driven by the tactile sensation, we reanalysed our main outcome parameters (MEP probability and RMT_Fit_ change) from experiments 2-4 after adding sensation (Exp. 2) or detection accuracy (Exp. 3-4) as a covariate (all covariates were z-scored due to non-normal distribution). After the covariates were added, all the effects reported in experiments 2-4 remained significant (all p ≤ 0.04) making it unlikely that tactile sensation was the main driver of our results. Furthermore, strength or accuracy of stimulation-induced sensations did not correlate with the measured effects, i.e., MEP probability and RMT_Fit_ change in experiments 2 and 3 (Exp.2: r = 0.22, p = 0.34; Exp.3: r = 0.04, p = 0.88). Only Exp.4 showed a significant correlation (r = 0.5, p = 0.02) which, however, could not have driven our results because better detection of tRNS diminished its effect on lowering RMT_Fit_. Similarly, there was also no significant correlation between the monitored impedance between the electrodes (which was always < 15 kΩ) and the observed effects related to tRNS (all r ≤ 0.21, p ≥ 0.4).

#### Background EMG

Analysis of bgEMG across the experimental conditions demonstrated that muscle activity was generally low (on average less than 0.0025 mV across experiments). Moreover, additional analyses using bgEMG as a covariate in experiments 2-4 revealed that all reported TMS effects remained significant (all p ≤ 0.03).

#### TRNS INDUCED EFFECTS DO NOT DEPEND ON MEP CRITERIA

The above results used MEP probability as the main outcome measurement. Importantly, the reported effects were not driven by our MEP amplitude criterion (i.e., MEP amplitude ≥ 0.05 mV), as control analyses with slightly different criteria (i.e., MEP cut-off amplitudes of 0.03-0.07 mV) revealed a similar pattern of results in all our experiments. Namely, irrespective of the adopted MEP criterion, we confirmed tRNS-induced enhancement in MEP probability in Exp. 1 (all r ≤ −0.56, z ≤ −2.86, p ≤ 0.004), we found a gradual increase in MEP probability for higher tRNS intensities in Exp. 2 (all F ≥ 2.5, p ≤ 0.05) and observed the decrease in RMT_Fit_ during 2mA tRNS over M1 (vs no tRNS in Exp. 3: all t ≥ 2.3, p ≤ 0.03 and vs 2mA tRNS over V1 in Exp. 4: all t ≥ 2.3, p ≤ 0.03).

#### TMS intensity adjustments

It is well known that the individual responsiveness to TMS can drift slightly throughout a TMS experiment due to state-dependent changes, which affect measurements close to RMT intensities in particular (Karabanov et al. 2015). As recommended by the Karabanov et al, we adjusted TMS intensities between experimental blocks to ensure subthreshold stimulation (i.e., p(MEP_0.05mV_) no tRNS ≤ 0.3 or ≤ 0.5) throughout Experiments 2, 3 and 4 as shown in **Figure 9**. Importantly, once TMS intensity was adjusted, we collected an equal amount of data for each of the tRNS conditions ensuring that the direct comparison of 2mA tRNS versus the control condition was not confounded by the between-block adjustments.

Next, we analysed whether the intensity adjustments followed a systematic trend by pooling these data across experiments and comparing the first to the last block. We found no systematic changes in TMS intensity (t_(63)_ = 1.15, p = 0.25, MD = 0.22 ± 1.5% **Figure 9**) making it unlikely that tRNS or other aspects of our experimental procedure caused any longer-lasting effects on M1 responsiveness.

#### Electric field measurement

Finally, we also tested electric field induced by TMS pulse with and without tRNS. The induced electric field was measured by oscilloscope using a search coil placed under the TMS coil. tRNS was delivered to a conducting phantom medium soaked in saline solution through two electrodes placed on either side of the coils (impedance = 2 kΩ). We measured 20 TMS pulses with and without electrical noise stimulation (2mA tRNS) in an alternating manner. We showed that tRNS did not influence the electric field induced by the TMS pulse (t_(19)_ = 0.6, p = 0.53, MD = 0.05 ± 0.3 μV). This confirmed that all changes in MEP probability originated from the modulation of M1 responsiveness and not stronger magnetic stimulation.

## Discussion

This study provides direct evidence that online tRNS over M1 enhances the responsiveness of cortical motor circuits via a shift in response threshold. Across four separate experiments we showed that acute tRNS effects manifested as (i) a higher probability of evoking MEPs when TMS was applied with intensities at or slightly below resting motor threshold (**Figure 4, Figure *5*, Figure *6*A, Figure *7*A**), and (ii) lower estimated resting motor thresholds (RMT_Fit_ shown in **Figure 6B-C, Figure *7*B-C**). Importantly, the observed effects appeared to be specific for tRNS delivery over M1 but not over V1 (**Figure 7B**), making it unlikely that our results were driven by any unspecific tRNS effects. Our findings consistently indicate that tRNS enhances the responsiveness of cortical motor circuits by lowering the response threshold and provide important proof-of-principle evidence of noise benefits at the neural population level in humans.

### acute tRNS-induced noise benefits are partly consistent with SR theory

We consistently observed that adding electrical noise to M1 can acutely increase its responsiveness to TMS. One potential explanation for the observed results is signal enhancement during acute noise delivery to a non-linear system. Such noise benefits are one hallmark feature of SR theory (McDonnell and Abbott 2009). A second important feature is that noise benefits are a function of noise intensity exhibiting an inverted U-shape dose-response relationship, i.e., too much noise is detrimental. While our neurophysiological study provided proof-of-principle evidence of noise benefits at the neural population level in human cortex, we did not demonstrate the inverted U-shape function for higher tRNS intensities (**Figure 5**) and can draw no inference about how much noise would be optimal for increasing the responsiveness of M1.

Previous studies (van der Groen and Wenderoth 2016; van der Groen et al. 2018; Pavan et al. 2019) demonstrated an inverted U-shape function of tRNS effects using behavioural outcome parameters that are typically acquired in accordance with detection theory. One explanation for the detrimental effect of excessive noise is that it causes the system to respond even if there is no signal to detect and reduces the detection rate by causing too many “*false alarms*”. However, in our experiments a *false alarm* would mean that tRNS alone would occasionally evoke MEPs. Thus, probing the detrimental effect of adding too much noise to the resting motor system via tRNS would require much higher intensities than used here, which would certainly cause strong discomfort in human participants (Fertonani et al. 2015).

Accordingly, we cannot claim that we added the optimal level of noise to M1 based on our data. In fact, the absolute changes in RMT were relatively small (even though statistical effect sizes were medium to large). This can be partly attributed to the fact that we aimed to modulate resting motor threshold, which is one of the most robust and reliable TMS measurements (Ngomo et al. 2012; Dissanayaka et al. 2018). Indeed, RMT has rarely been modulated by other forms of electrical stimulation (Nitsche et al. 2005), including long-lasting effects of tRNS (Terney et al. 2008). However, another explanation for the relatively small absolute effects is that tRNS induced a suboptimal amount of noise.

### Alternative accounts of the observed acute tRNS-induced noise benefits

Until now, a change in corticomotor excitability measurements has been demonstrated after prolonged tRNS delivery and has been hypothesized to reflect long-lasting neuroplastic changes (Terney et al. 2008; Chaieb et al. 2011, 2015; Abe et al. 2019; Moret et al. 2019). However, neuroplastic changes are unlikely to have driven the acute tRNS effects in our study since tRNS conditions were always interleaved with no noise or other control conditions, thereby minimizing the influence of long-term excitability changes. Moreover, there was no systematic change in adjusted TMS intensity for the no tRNS condition over time, making it unlikely that our results were affected by long-term neuroplasticity (**Figure 9**).

Another possible mechanism is that repeated subthreshold stimulations with tRNS induced consecutive openings of sodium channels which might lead to temporal summation of small membrane potentials, cause depolarization of the neural membrane due to an increased influx of inward sodium currents and/or prevent the homeostasis of the system (Terney et al. 2008; Fertonani et al. 2011). This may affect excitability of M1 in a similar manner as reported here.

Finally, the effects of tRNS may also be attributed to the increased synchronization of neural firing through amplification of subthreshold oscillatory activity, reducing the amount of endogenous noise (Miniussi et al. 2013).

### Possible neurophysiological substrate mediating tRNS-induced noise benefits

We showed that a short tRNS bout of less than 3 seconds acutely influences the neurophysiology of human motor cortex. Using threshold estimation analysis as our primary outcome measurement (Vucic et al. 2018), we showed that tRNS specifically affected the response threshold of cortical motor circuits. By contrast, tRNS had no systematic effect on MEP amplitude. It has been argued that motor threshold and MEP amplitude reflect independent neural processes (Paulus et al. 2008; Vucic and Kiernan 2017; Vucic et al. 2018).

From a neurophysiological point of view, the motor threshold reflects the efficacy of a chain of synapses from cortical interneurons in layer II/III to the muscles (Kobayashi and Pascual-Leone 2003). Pharmacological studies have implicated voltage-gated sodium channels as a major determinant of motor threshold since blocking sodium channels increases RMT (Tergau et al. 2003; Sommer et al. 2012; Ziemann et al. 2015).

Even though the neurophysiological mechanism of tRNS is not completely understood, previous studies (Onorato et al. 2016) suggested that such enhancing a neurons response via electrical noise occurred by the concurrent activation of voltage-gated sodium channels. A recent study measured sodium currents in somatosensory and auditory pyramidal neurons *in vitro* while stimulating the cells with different levels of electrical random noise (Remedios et al. 2019). The authors showed that some neurons exhibited higher peak amplitudes of sodium currents, elicited by a voltage-clamp-ramp protocol, which correlated with shorter latencies of the neuronal response during brief electrical noise delivery. In other words, in those neurons, less voltage was needed to evoke the sodium current peak when the optimal level of electrical noise was delivered. Additionally, tRNS might affect neuronal populations by increasing the probability of synchronized firing within neuronal populations (Miniussi et al. 2013). In this regard, it has been estimated that alternating currents at 100 Hz polarize a single neuron by a relatively small amount (Deans et al. 2007). However, networks of many synaptically connected active neurons reveal higher sensitivity to field modulation than single cells, thus, amplifying the stimulation effect (Fröhlich and McCormick 2010; Reato et al. 2010). Accordingly, even subthreshold stimulation that induces very weak electric fields in cortex can modulate membrane potentials (Gluckman et al. 1996; Francis et al. 2003; Bikson et al. 2006). Based on these findings we propose that tRNS might have reduced RMT_Fit_ by slightly increasing the responsiveness of voltage-gated sodium channels (Ziemann et al. 2015) in large populations of cortical cells.

Interestingly, unlike short 4s-bouts of tDCS (Nitsche et al., 2003; but see example for large variability of tDCS effects on MEP amplitude Jonker et al., 2021), acute tRNS induced no changes in MEP amplitudes. In line with this finding and the supposed role of sodium channels, previous pharmacological studies have shown that blocking voltage-gated sodium channels had no or only inconsistent effects on MEP amplitudes (Paulus et al. 2008; Vucic et al. 2018).

### Increased cortical responsiveness via tRNS is unlikely to result from tactile stimulation

In our study, many participants felt a slight but noticeable skin sensation (Fertonani et al. 2015). This constitutes a potential confound because participants were not blinded to tRNS conditions and because some effects might be driven by transcutaneous stimulation of peripheral nerves rather than by transcranial stimulation of cortical neurons (Asamoah et al. 2019). We reanalysed all our experiments accounting for tactile sensation and showed that they did not contribute to the observed effects. Most importantly, in the final experiment we utilized an active control condition. Behavioural results revealed that the intensity of the skin sensation was similar but only M1 stimulation lowered the response threshold of cortical motor circuits (**Figure 7**). Moreover, a study in non-human primates found significant neuromodulation effects despite blocking or substantially suppressing somatosensory input (Vieira et al. 2020). Thus, we argue that the effects observed in our study are most likely caused by adding electrical noise to M1 rather than by unspecific effects of tRNS.

### Summary and conclusion

Our results demonstrate that tRNS changes the responsiveness of M1 circuits. We observed acute modulation of RMT_Fit_, which seems to reflect immediate signal enhancement rather than neuroplastic changes. Such increase in responsiveness of a nonlinear system in the presence of noise seems to be consistent with one of the two hallmarks of SR theory showing noise benefits at the neural population level in humans (McDonnell and Abbott 2009). Our study provides evidence that online tRNS influences the neurophysiology of the human motor cortex and sheds new light on understanding the impact of acute electrical noise stimulation on neural processing at the network level.

## Notes

We thank all the participants for their time and effort, Daniel Woolley for his valuable help and feedback on the manuscript, Ernest Mihelj for his contribution to the data analysis, and Marta Stepien, Lena Salzmann, Alexandra Burgler, and Luana Giacone for their assistance during data collection.

## Supplementary Material

Supplementary material is available at bioRxiv online.

## REFERENCES

Abe T, Miyaguchi S, Otsuru N, Onishi H. 2019. The effect of transcranial random noise stimulation on corticospinal excitability and motor performance. Neurosci Lett. 705:138– 142.

Antal A, Herrmann CS. 2016. Transcranial Alternating Current and Random Noise Stimulation: Possible Mechanisms. Neural Plast. 2016:1–12.

Asamoah B, Khatoun A, Mc Laughlin M. 2019. tACS motor system effects can be caused by transcutaneous stimulation of peripheral nerves. Nat Commun. 10:1–16.

Bikson M, Radman T, Datta A. 2006. Rational modulation of neuronal processing with applied electric fields. 2006 Int Conf IEEE Eng Med Biol Soc. 1616–1619.

Chaieb L, Antal A, Paulus W. 2015. Transcranial random noise stimulation-induced plasticity is NMDA-receptor independent but sodium-channel blocker and benzodiazepines sensitive. Front Neurosci. 9:1–9.

Chaieb L, Paulus W, Antal A. 2011. Evaluating aftereffects of short-duration transcranial random noise stimulation on cortical excitability. Neural Plast. 2011.

Davila-Pérez P, Jannati A, Fried PJ, Cudeiro Mazaira J, Pascual-Leone A. 2018. The Effects of Waveform and Current Direction on the Efficacy and Test–Retest Reliability of Transcranial Magnetic Stimulation. Neuroscience. 393:97–109.

Deans JK, Powell AD, Jefferys JGR. 2007. Sensitivity of coherent oscillations in rat hippocampus to AC electric fields. J Physiol. 583:555–565.

Dissanayaka T, Zoghi M, Farrell M, Egan G, Jaberzadeh S. 2018. Comparison of Rossini– Rothwell and adaptive threshold-hunting methods on the stability of TMS induced motor evoked potentials amplitudes. J Neurosci Res. 96:1758–1765.

Faul F, Erdfelder E, Lang A, Buchner A. 2007. G * Power 3?: A flexible statistical power analysis program for the social, behavioral, and biomedical sciences. Behav Res Methods. 39:175–191.

Fertonani A, Ferrari C, Miniussi C. 2015. What do you feel if I apply transcranial electric stimulation? Safety, sensations and secondary induced effects. Clin Neurophysiol. 126:2181–2188.

Fertonani A, Pirulli C, Miniussi C. 2011. Random Noise Stimulation Improves Neuroplasticity in Perceptual Learning. J Neurosci. 31:15416–15423.

Francis JT, Gluckman BJ, Schiff SJ. 2003. Sensitivity of Neurons to Weak Electric Fields. J Neurosci. 23:7255–7261.

Fröhlich F, McCormick DA. 2010. Endogenous electric fields may guide neocortical network activity. Neuron. 67:129–143.

Ghin F, Pavan A, Contillo A, Mather G. 2018. The effects of high-frequency transcranial random noise stimulation (hf-tRNS) on global motion processing?: An equivalent noise approach. Brain Stimul. 11:1263–1275.

Gingl Z, Kiss LB, Moss F. 1995. Non-dynamical stochastic resonance: Theory and experiments with white and arbitrarily coloured noise. Europhys Lett. 29:191–196.

Gluckman BJ, Spano ML, Netoff TI, Neel EJ, Schiff SJ, Spano WL. 1996. Stochastic Resonance in a Neuronal Network from Mammalian Brain. Phys Rev Lett. 77:4098–4101.

Groen O Van Der, Wenderoth N. 2016. Externally applied noise influences state-switching dynamics of binocular rivalry. 36:1002991.

Hallett M. 2007. Transcranial magnetic stimulation: A primer. Neuron. 55:187–199.

Herpich F, Melnick M, Agosta S, Huxlin K, Tadin D, Battelli L. 2019. Boosting learning efficacy with non-invasive brain stimulation in intact and brain-damaged humans. J Neurosci. 39:5551–5561.

Hinder MR, Goss EL, Fujiyama H, Canty AJ, Garry MI, Rodger J, Summers JJ. 2014. Inter- and intra-individual variability following intermittent theta burst stimulation: Implications for rehabilitation and recovery. Brain Stimul. 7:365–371.

Jannati A, Fried PJ, Block G, Oberman LM, Rotenberg A, Pascual-Leone A. 2019. Test– Retest Reliability of the Effects of Continuous Theta-Burst Stimulation. Front Neurosci. 13:447.

Jonker ZD, Gaiser C, Tulen JHM, Ribbers GM, Frens MA, Selles RW. 2021. No effect of anodal tDCS on motor cortical excitability and no evidence for responders in a large double-blind placebo-controlled trial. Brain Stimul. 14:100–109.

Karabanov AN, Raffin E, Siebner HR. 2015. The resting motor threshold - Restless or resting? A repeated threshold hunting technique to track dynamic changes in resting motor threshold. Brain Stimul. 8:1191–1194.

Kobayashi M, Pascual-Leone A. 2003. Transcranial magnetic stimulation in neurology. Lancet Neurol. 2:145–156.

Lakens D. 2013. Calculating and reporting effect sizes to facilitate cumulative science: a practical primer for t-tests and ANOVAs. Front Psychol. 4:1–12.

Livingston SC, Ingersoll CD. 2008. Intra-rater reliability of a transcranial magnetic stimulation technique to obtain motor evoked potentials. Int J Neurosci. 118:239–256.

McDonnell MD, Abbott D. 2009. What is stochastic resonance? Definitions, misconceptions, debates, and its relevance to biology. PLoS Comput Biol. 5.

Mills KR, Boniface SJ, Schubert M. 1992. Magnetic brain stimulation with a double coil: the importance of coil orientation. Electroencephalogr Clin Neurophysiol Evoked Potentials. 85:17–21.

Miniussi C, Harris JA, Ruzzoli M. 2013. Modelling non-invasive brain stimulation in cognitive neuroscience. Neurosci Biobehav Rev. 37:1702–1712.

Moliadze V, Atalay D, Antal A, Paulus W. 2012. Close to threshold transcranial electrical stimulation preferentially activates inhibitory networks before switching to excitation with higher intensities. Brain Stimul. 5:505–511.

Moret B, Camilleri R, Pavan A, Lo Giudice G, Veronese A, Rizzo R, Campana G. 2018. Differential effects of high-frequency transcranial random noise stimulation (hf-tRNS) on contrast sensitivity and visual acuity when combined with a short perceptual training in adults with amblyopia. Neuropsychologia. 114:125–133.

Moret B, Donato R, Nucci M, Cona G, Campana G. 2019. Transcranial random noise stimulation (tRNS): a wide range of frequencies is needed for increasing cortical excitability. Sci Rep. 9:1–8.

Moss F, Ward LM, Sannita WG. 2004. Stochastic resonance and sensory information processing: A tutorial and review of application. Clin Neurophysiol. 115:267–281.

Ngomo S, Leonard G, Moffet H, Mercier C. 2012. Comparison of transcranial magnetic stimulation measures obtained at rest and under active conditions and their reliability. J Neurosci Methods. 205:65–71.

Nitsche MA, Cohen LG, Wassermann EM, Priori A, Lang N, Antal A, Paulus W, Hummel F, Boggio PS, Fregni F, Pascual-Leone A. 2008. Transcranial direct current stimulation: State of the art 2008. Brain Stimul. 1:206–223.

Nitsche MA, Fricke K, Henschke U, Schlitterlau A, Liebetanz D, Lang N, Henning S, Tergau F, Paulus W. 2003. Pharmacological modulation of cortical excitability shifts induced by transcranial direct current stimulation in humans. J Physiol. 293–301.

Nitsche MA, Seeber A, Frommann K, Klein CC, Rochford C, Nitsche MS, Fricke K, Liebetanz D, Lang N, Antal A, Paulus W, Tergau F. 2005. Modulating parameters of excitability during and after transcranial direct current stimulation of the human motor cortex. J Physiol. 568:291–303.

Oldfield RC. 1971. The Assesment and Analysis of Handedness: The Edinburgh Inventory. 9:97–113.

Onorato I, D’Alessandro G, Di Castro MA, Renzi M, Dobrowolny G, Musarò A, Salvetti M, Limatola C, Crisanti A, Grassi F. 2016. Noise enhances action potential generation in mouse sensory neurons via stochastic resonance. PLoS One. 11:1–12.

Paulus W, Classen J, Cohen LG, Large CH, Di Lazzaro V, Nitsche M, Pascual-Leone A, Rosenow F, Rothwell JC, Ziemann U. 2008. State of the art: Pharmacologic effects on cortical excitability measures tested by transcranial magnetic stimulation. Brain Stimul. 1:151–163.

Pavan A, Ghin F, Contillo A, Milesi C, Campana G, Mather G. 2019. Modulatory mechanisms underlying high-frequency transcranial random noise stimulation (hf-tRNS): A combined stochastic resonance and equivalent noise approach. Brain Stimul. 12:967–977.

Pearson K. 1897. Mathematical contributions to the theory of evolution.—on a form of spurious correlation which may arise when indices are used in the measurement of organs. Proc R Soc london. 60:489–498.

Pirulli C, Fertonani A, Miniussi C. 2013. The role of timing in the induction of neuromodulation in perceptual learning by transcranial electric stimulation. Brain Stimul. 6:683–689.

Pirulli C, Fertonani A, Miniussi C. 2016. On the Functional Equivalence of Electrodes in Transcranial Random Noise Stimulation. Brain Stimul. 9:621–622.

Rathelot J-A, Strick PL. 2009. Subdivisions of primary motor cortex based on cortico-motoneuronal cells. Proc Natl Acad Sci. 106:918–923.

Reato D, Rahman A, Bikson M, Parra LC. 2010. Low-Intensity Electrical Stimulation Affects Network Dynamics by Modulating Population Rate and Spike Timing. J Neurosci. 30:15067–15079.

Remedios L, Mabil P, Flores-Hernández J, Torres-Ramírez O, Huidobro N, Castro G, Cervantes L, Tapia JA, De la Torre Valdovinos B, Manjarrez E. 2019. Effects of Short-Term Random Noise Electrical Stimulation on Dissociated Pyramidal Neurons from the Cerebral Cortex. Neuroscience. 404:371–386.

Rossi S, Hallett M, Rossini PM, Pascual-Leone A, Avanzini G, Bestmann S, Berardelli A, Brewer C, Canli T, Cantello R, Chen R, Classen J, Demitrack M, Di Lazzaro V, Epstein CM, George MS, Fregni F, Ilmoniemi R, Jalinous R, Karp B, Lefaucheur JP, Lisanby S, Meunier S, Miniussi C, Miranda P, Padberg F, Paulus W, Peterchev A, Porteri C, Provost M, Quartarone A, Rotenberg A, Rothwell J, Ruohonen J, Siebner H, Thut G, Valls-Solè J, Walsh V, Ugawa Y, Zangen A, Ziemann U. 2009. Safety, ethical considerations, and application guidelines for the use of transcranial magnetic stimulation in clinical practice and research. Clin Neurophysiol. 120:2008–2039.

Rossini P, Barker A, Berardelli A. 1994. Non-invasive electrical and magnetic stimulation of the brain, spinal cord and roots: basic principles and procedures for routine clinical application. Report of an IFCN. Electroencephalogr Clin Neurophysiol. 91:79–92.

Schambra HM, Ogden RT, Martínez-Hernández IE, Lin X, Chang YB, Rahman A, Edwards DJ, Krakauer JW. 2015. The reliability of repeated TMS measures in older adults and in patients with subacute and chronic stroke. Front Cell Neurosci. 9:1–18.

Simonotto E, Riani M, Seife C, Roberts M, Twitty J, Moss F. 1997. Visual Perception of Stochastic Resonance. 256:6–9.

Sommer M, Gileles E, Knappmeyer K, Rothkegel H, Polania R, Paulus W. 2012. Carbamazepine reduces short-interval interhemispheric inhibition in healthy humans. Clin Neurophysiol. 123:351–357.

Tergau F, Wischer S, Somal HS, Nitsche MA, Mercer AJ, Paulus W, Steinhoff BJ. 2003. Relationship between lamotrigine oral dose, serum level and its inhibitory effect on CNS: Insights from transcranial magnetic stimulation. Epilepsy Res. 56:67–77.

Terney D, Chaieb L, Moliadze V, Antal A, Paulus W. 2008. Increasing Human Brain Excitability by Transcranial High-Frequency Random Noise Stimulation. J Neurosci. 28:14147– 14155.

Thielscher A, Antunes A, Saturnino GB. 2015. Field modeling for transcranial magnetic stimulation: A useful tool to understand the physiological effects of TMS? Proc Annu Int Conf IEEE Eng Med Biol Soc EMBS. 222–225.

Thielscher A, Saturnino GB. 2019. Field calculations with SimNIBS. In: Brainbox Initiative Conference. Tu YK. 2016. Testing the relation between percentage change and baseline value. Sci Rep. 6:23247.

van der Groen O, Mattingley JB, Wenderoth N. 2019. Altering brain dynamics with transcranial random noise stimulation. Sci Rep. 9:1–8.

van der Groen O, Tang MF, Wenderoth N, Mattingley JB. 2018. Stochastic resonance enhances the rate of evidence accumulation during combined brain stimulation and perceptual decision-making. PLoS Comput Biol. 14:1–17.

van der Groen O, Wenderoth N. 2016. Transcranial Random Noise Stimulation of Visual Cortex: Stochastic Resonance Enhances Central Mechanisms of Perception. J Neurosci. 36:5289–5298.

Vieira PG, Krause MR, Pack CC. 2020. tACS entrains neural activity while somatosensory input is blocked. PLoS Biol. 18:1–14.

Vöröslakos M, Takeuchi Y, Brinyiczki K, Zombori T, Oliva A, Fernández-Ruiz A, Kozák G, Kincses ZT, Iványi B, Buzsáki G, Berényi A. 2018. Direct effects of transcranial electric stimulation on brain circuits in rats and humans. Nat Commun. 9.

Vucic S, Kiernan MC. 2017. Transcranial Magnetic Stimulation for the Assessment of Neurodegenerative Disease. Neurotherapeutics. 14:91–106.

Vucic S, van den Bos M, Parvathi M, Howells J, Dharmadasa T, Kiernan MC. 2018. Utility of threshold tracking transcranial magnetic stimulation in ALS. Clin Neurophysiol Pract. 3:164–172.

Wassermann EM. 1998. Risk and safety of repetitive transcranial magnetic stimulation. Electroencephalogr Clin Neurophysiol. 108:1–16.

Ziemann U, Reis J, Schwenkreis P, Rosanova M, Strafella A, Badawy R, Müller-Dahlhaus F. 2015. TMS and drugs revisited 2014. Clin Neurophysiol. 126:1847–1868.

