## SUPPLEMENTARY MATERIAL for "Transcranial Random Noise Stimulation acutely lowers the response threshold of human motor circuits"

##### MEP AMPLITUDE

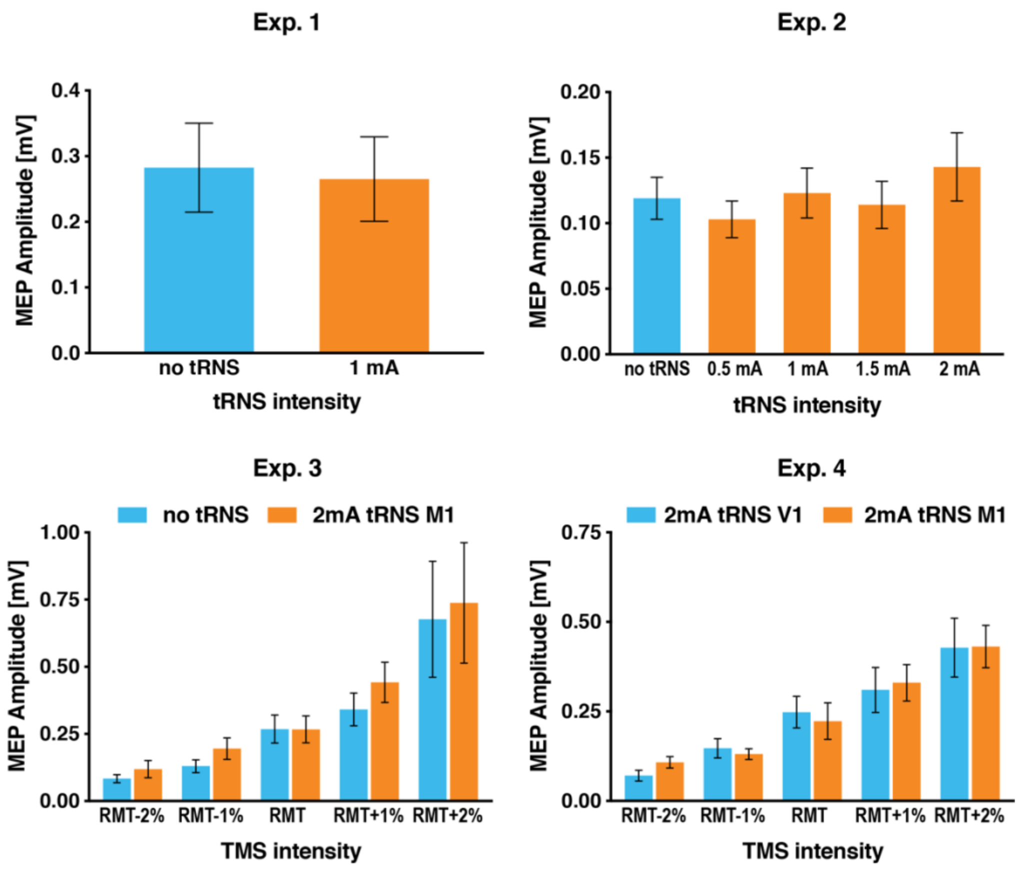


**Figure S1** Average motor evoked potential (MEP) amplitude elicited by single-pulse transcranial magnetic stimulation (TMS) in the experimental and control conditions. tRNS modulated MEP amplitude to a minor extent resulting in significant effects only in Exp. 3 ($p = 0.005$), but not in Exp. 1, 2 and 4 ($p \geq 0.23$). Error bars indicate SE.

#### TACTILE SENSATION

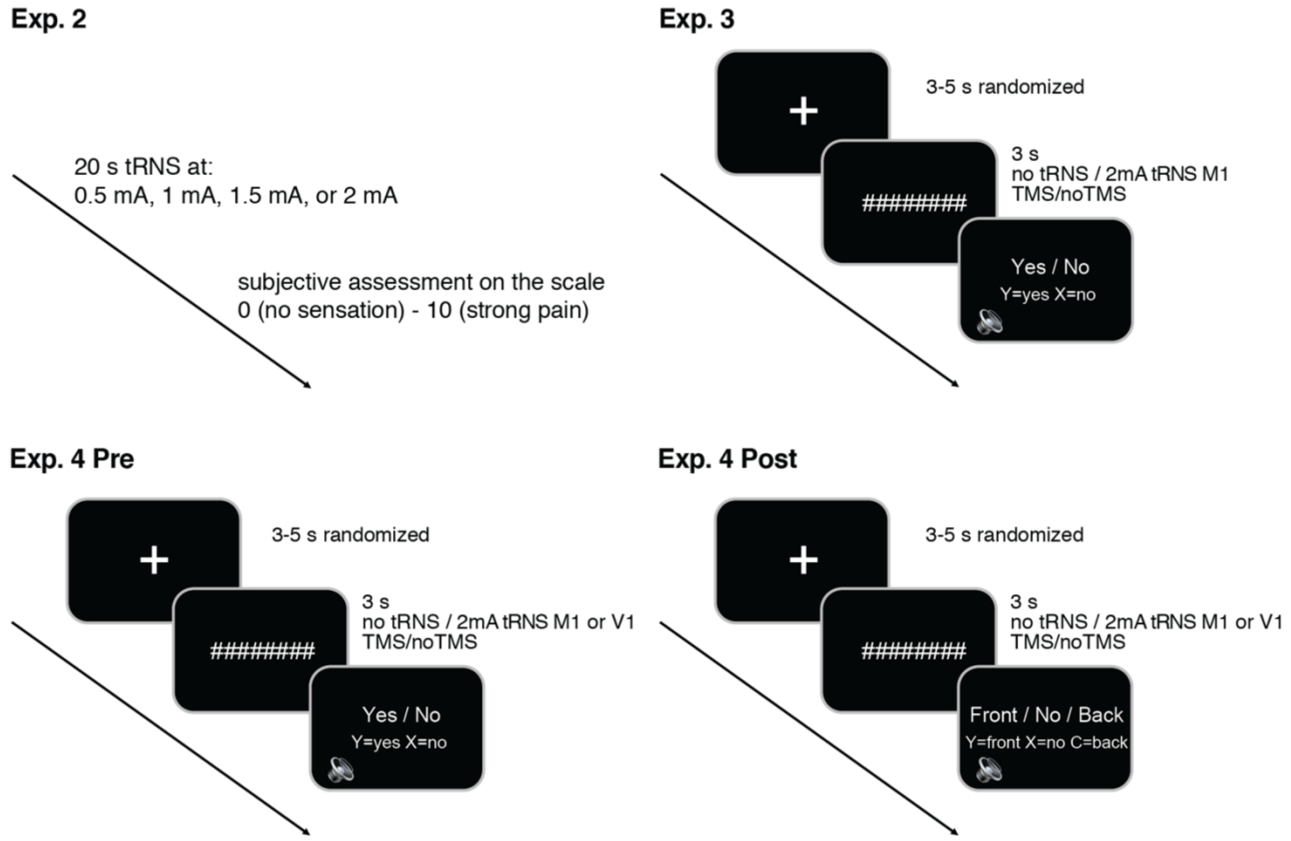


**Figure S2** Tactile sensation assessment in experiments 2-4. **Exp. 2:** Before the experiment all experimental transcranial random noise stimulation (tRNS) intensities (0.5-2mA) were presented to the participant for 20 s in a randomized order to make sure that the stimulation did not cause any unpleasant sensations. Participants were asked to subjectively assess the strength of tRNS on a scale from 0 (no sensation) to 10 (pain). **Exp. 3:** The detection task consisted of 20 trials. Participants received either tRNS (2mA, with and without transcranial magnetic stimulation (TMS) on half of the trials) or no tRNS (with and without TMS). Their task on each trial was to indicate (after an auditory cue) if they felt something underneath the tRNS electrodes (ignoring TMS pulses) by pressing the appropriate button on a keyboard. **Exp. 4 Pre:** The task was similar to Exp. 3, but with three stimulation conditions: no tRNS, 2mA tRNS over primary motor cortex (M1) or 2mA tRNS over primary visual cortex (V1). **Exp. 4 Post:** The task was similar to Exp. 4 Pre, but here participants were asked to differentiate between 3 possible scenarios: stimulation located toward the front of the head (2mA tRNS over M1), toward the back of the head (2mA tRNS over V1), or no stimulation (no tRNS). The subjectively assessed sensation (Exp. 2) and detection accuracy (Exp. 3-4) were non-normal distributed so we normalized them using z-score before including into further analyses as covariates.

#### tRNS INDUCED EFFECTS DO NOT DEPEND ON MEP CRITERIA

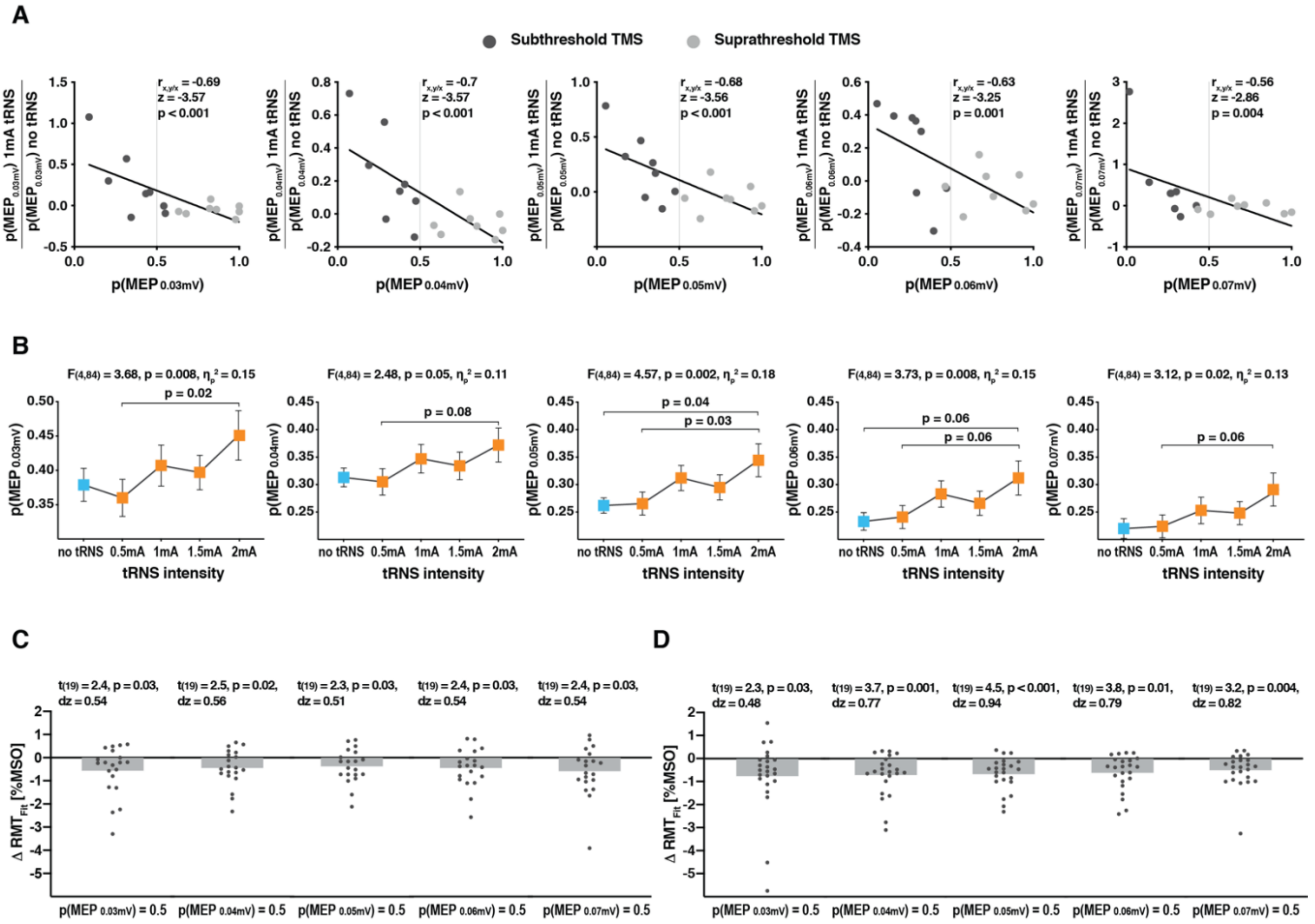


**Figure S3** Control analyses with different motor evoked potential amplitude criteria (i.e. MEP cut-off amplitudes of 0.03-0.07 mV) revealed a similar pattern of results in all our experiments. **A** Experiment 1: Significant increase in MEP probability (p(MEP)) during transcranial random noise stimulation (tRNS) delivery for subthreshold transcranial magnetic stimulation (TMS) regardless of the MEP criterion. **B** Experiment 2: Probability of MEP elicited by subthreshold TMS at different tRNS intensities applied over primary motor cortex (M1). Regardless of the MEP criterion the results reveal a gradually increasing pattern of MEP probability for higher tRNS intensities, with the 2mA stimulation being the most effective. Error bars indicate SE. **C** Experiment 3: Modulation of rest motor threshold (RMT): individual differences between $RMT_{Fit}$ in the 2mA tRNS over M1 and no tRNS condition. Most of the participants demonstrated a decrease in the $RMT_{Fit}$ for 2mA tRNS over M1 vs no tRNS control condition regardless of the MEP criterion. **D** Experiment 4: Modulation of RMT: individual differences between $RMT_{Fit}$ in 2mA tRNS over M1 vs control stimulation site. The majority of participants demonstrated a decrease in $RMT_{Fit}$ for tRNS over M1 in comparison to the control site stimulation condition regardless of the MEP criterion.

### ELECTRIC FIELD MEASUREMENT

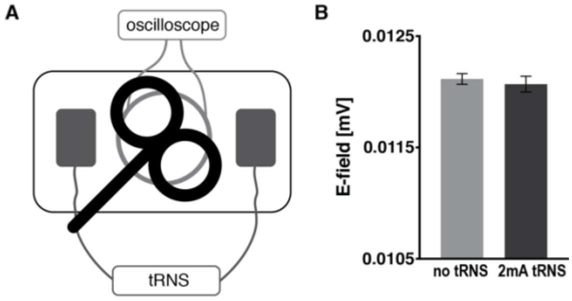


**Figure S4** Electric field induced by TMS pulse with and without tRNS. **A** Experimental set-up. The induced electric field was measured by oscilloscope using a search coil placed under the TMS coil. tRNS was delivered to a conducting phantom medium soaked in saline solution through two electrodes placed on either side of the coils (impedance = 2 kΩ). We measured 20 TMS pulses with and without electrical noise stimulation (2mA tRNS) in an alternating manner. **B** No difference between the two conditions was found suggesting that the obtained effects during the noise stimulation of left M1 originated from the modulation of motor cortex responsiveness, and not a modulated TMS pulse. Error bars indicate SE.
